# Cell cycle-independent integration of stress signals promotes Non-G1/G0 quiescence entry

**DOI:** 10.1101/2021.03.13.434817

**Authors:** Orlando Argüello-Miranda, Ashley Marchand, Taylor Kennedy, Marielle AX Russo, Jungsik Noh

## Abstract

Cellular quiescence is a non-proliferative state required for cell survival under stress and during development. In most quiescent cells, proliferation is stopped in a reversible state of low Cdk1 kinase activity; in many organisms, however, quiescent states with high Cdk1 activity can also be established through still uncharacterized stress or developmental mechanisms. Here, we used a microfluidics approach coupled to phenotypic classification by machine learning to identify stress pathways associated with starvation-triggered high-Cdk1 quiescent states in *Saccharomyces cerevisiae*. We found that low- and high-Cdk1 quiescent states shared a core of stress-associated processes, such as autophagy, protein aggregation, and mitochondrial upregulation, but differed in the nuclear accumulation of the stress transcription factors Xbp1, Gln3, and Sfp1. The decision between low- or high-Cdk1 quiescence was controlled by cell cycle-independent accumulation of Xbp1, which acted as a time-delayed integrator of the duration of stress stimuli. Our results show how cell cycle-independent stress-activated factors promote cellular quiescence outside of G1/G0.

## Introduction

Quiescence is a non-proliferative cellular state regulated by stimuli including starvation, DNA damage, and developmental signals (Cheung and Rando, 2013, van Velthoven and Rando, 2019). In unicellular organisms, quiescent states are essential for surviving adverse environmental conditions ranging from starvation to antibiotic exposure (Rittershaus et al., 2013). In multicellular organisms, quiescence is essential for tissue homeostasis and the maintenance of adult stem cells (Sun and Buttitta, 2017, Bainbridge, 2013, Apte et al., 2015, Nakamura-Ishizu et al., 2014, Li and Clevers, 2010).

Quiescence in eukaryotes commonly occurs after cells halt proliferation in a state where the activity of the proliferation-promoting kinase Cdk1 is low (low-Cdk1 quiescence). In many organisms, however, cells can enter quiescence in states of high-Cdk1 activity (high-Cdk1 quiescence), which are essential for development and single-cell survival (Wei et al., 1993, Hajeri et al., 2005, Velappan et al., 2017, Baisch, 1988). High-Cdk1 quiescence occurs in mammalian oocyte development (Nixon et al., 2002), embryonic development of invertebrates (Hajeri et al., 2005, Nystul et al., 2003, Otsuki and Brand, 2018), plant meristems (Velappan et al., 2017), microorganisms (Klosinska et al., 2011) and metastasis-initiating cancer cells (Wang et al., 2015). Despite its ubiquitous nature, mechanisms for high-Cdk1 quiescence remain largely uncharacterized (Sun and Gresham, 2020).

Several conserved mechanisms maintain low-Cdk1 quiescence including: (1) the accumulation of Cdk1 transcriptional repressors, such as Whi5/Rb (2) the accumulation of Cdk1 inhibitors, such as Sic1/p27/p21, and (3) the destruction of Cdk1 activators by APC/C-Cdh1-mediated proteolysis (Moser et al., 2018, Cappell et al., 2016, Hopkins et al., 2017). By contrast, only few factors promoting high-Cdk1 quiescence have been identified, such as *C. elegans* spindle assembly components SAN-1 and MDF-2 (Nystul et al., 2003) or the pseudokinase Tribbles (Trbl) in *D. melanogaster*’s neural stem cells (Otsuki and Brand, 2018).

Stress-activated pathways are good candidates to promote quiescence regardless of Cdk1 activity due to their capacity to decrease overall translation, transcription, and modulate cell cycle protein networks (Miles et al., 2013, Marion et al., 2004, Escote et al., 2004, Wysocki et al., 2006). However, quiescence-promoting stimuli, such as starvation, often trigger multiple stress signaling pathways that are experimentally challenging to simultaneously measure in single cells (De Virgilio, 2012) and high-Cdk1 quiescent states are particularly difficult to characterize because they are less common than low-Cdk1 quiescence or occur in complex tissues (Velappan et al., 2017, Otsuki and Brand, 2018). These challenges can be overcome by studying quiescence in a tractable unicellular model organism such as *Saccharomyces cerevisiae;* however, most protocols to induce quiescence require long-term batch cultures, where interactions between cells and changing chemical and physical medium parameters confound the assessment of whether stress factors are associated or causative of quiescent states (Klosinska et al., 2011). Furthermore, because single cells are not tracked in batch cultures, it remains controversial whether high-Cdk1 quiescent states in S. *cerevisiae* correspond to a failure of arresting cell cycle in response to starvation, a population of slowly dividing cells, or a proper cellular strategy to survive acute starvation stress after G1 exit.

In this work, we used a microfluidics assay with controlled environmental conditions to study S. *cerevisiae’s* transition from proliferation into quiescence after acute starvation. To characterize quiescent states with high-Cdk1 activity, we used machine learning (ML) algorithms to automatically distinguish proper high-Cdk1 quiescent states from slowly dividing or senescent cells. Quiescent states were classified based on established markers of the high-Cdk1 activity state, such as destruction of the Cdk1 inhibitor Sic1, inactivation of APC/C-Cdh1-mediated proteolysis, and assembly of the septin ring at the budding site after G1 exit (Zhang et al., 2011).

We found that high-Cdk1 quiescent states reproducibly occurred when a proliferating population was challenged by acute starvation. The frequency of high-Cdk1 quiescent cells depended on the stress status of the population at starvation onset. Although all quiescent states shared a core of stress-associated processes such as the upregulation of autophagy, protein aggregation, and mitochondrial biomass, high-Cdk1 quiescence was distinguished by increased nuclear accumulation of the general transcriptional repressor Xbp1, the nitrogen stress transcription factor Gln3, and the ribosome biogenesis factor Sfp1. The establishment of high-Cdk1 quiescence was controlled by cell cycle-independent nuclear accumulation of Xbp1, which acted as a time-delayed integrator of stress stimuli. Our results show that cell cycle independent integration of stress stimuli by transcriptional repressors is a viable cellular strategy to establish quiescence outside of a G1/G0 state.

## Results

### Acute starvation induces heterogeneous quiescent states

To study the establishment of quiescence in single cells, we modified a previously published microfluidics assay for quiescence induction in *S. cerevisiae* (Arguello-Miranda et al., 2018). Briefly, proliferating W303 diploid cells were cultured in a flow cell with rich medium (synthetic complete defined, SCD) before being exposed to 20 h of acute starvation medium (0.7% potassium acetate, pH 7.2, 0.6 psi, which is a modified version of meiosis-inducing medium (Buonomo et al., 2000)). Upon acute starvation, cells recapitulated quiescence properties previously observed in batch cultures (Miles et al., 2019). Proliferation was stopped after a slow and asymmetric final cell division, which required 5 ± 3 h under starvation conditions as opposed to 93 ± 7 min in SCD (Fig. 1 A). The daughter/mother cell size ratio at cytokinesis decreased from 0.58 ± 0.5 in rich medium to 0.25 ± 0.04 for cells born after starvation (Fig. 1 B, C and D). As reported in quiescent batch cultures (Shi et al., 2010), cells upregulated trehalose and glycogen metabolism during quiescence entry as measured by tracking the level of the mNeonGreen (mNG)-tagged trehalose synthase catalytic subunit, Tps1, and the mNG-tagged glycogen synthase, Gsy2 (Fig. 1 E). Quiescent cells (Q-cells) maintained high viability as judged by return to proliferation (Fig. 1 F, 72 h is the limit of the microfluidics device) and acquired a stress-resistant state after 20 h of starvation as judged by 94 ± 3% survival (n=7, 443 cells) after exposure to 4 M NaCl for 4 h (Video 1). Strikingly, although most cells arrested in G1, as judged by maintaining an unbudded state, a reproducible minority of 7 ± 3 % cells remained arrested in a budded state (Fig. 1, G and H), which was highly viable upon return to rich medium (Fig. 1 I). Single-cell tracking of budded and unbudded quiescent cells showed no significant differences in cell volume (Fig. 1 J), Tps1 or Gsy2 upregulation (Fig. 1 K and L), or cell morphology (Fig. S1 A), as assessed by the overlap of the 95 % confidence intervals. We concluded that a fraction of proliferating S. *cerevisiae* cells entered viable quiescent states outside G1/G0 when challenged by acute starvation.

**Fig. 1.**
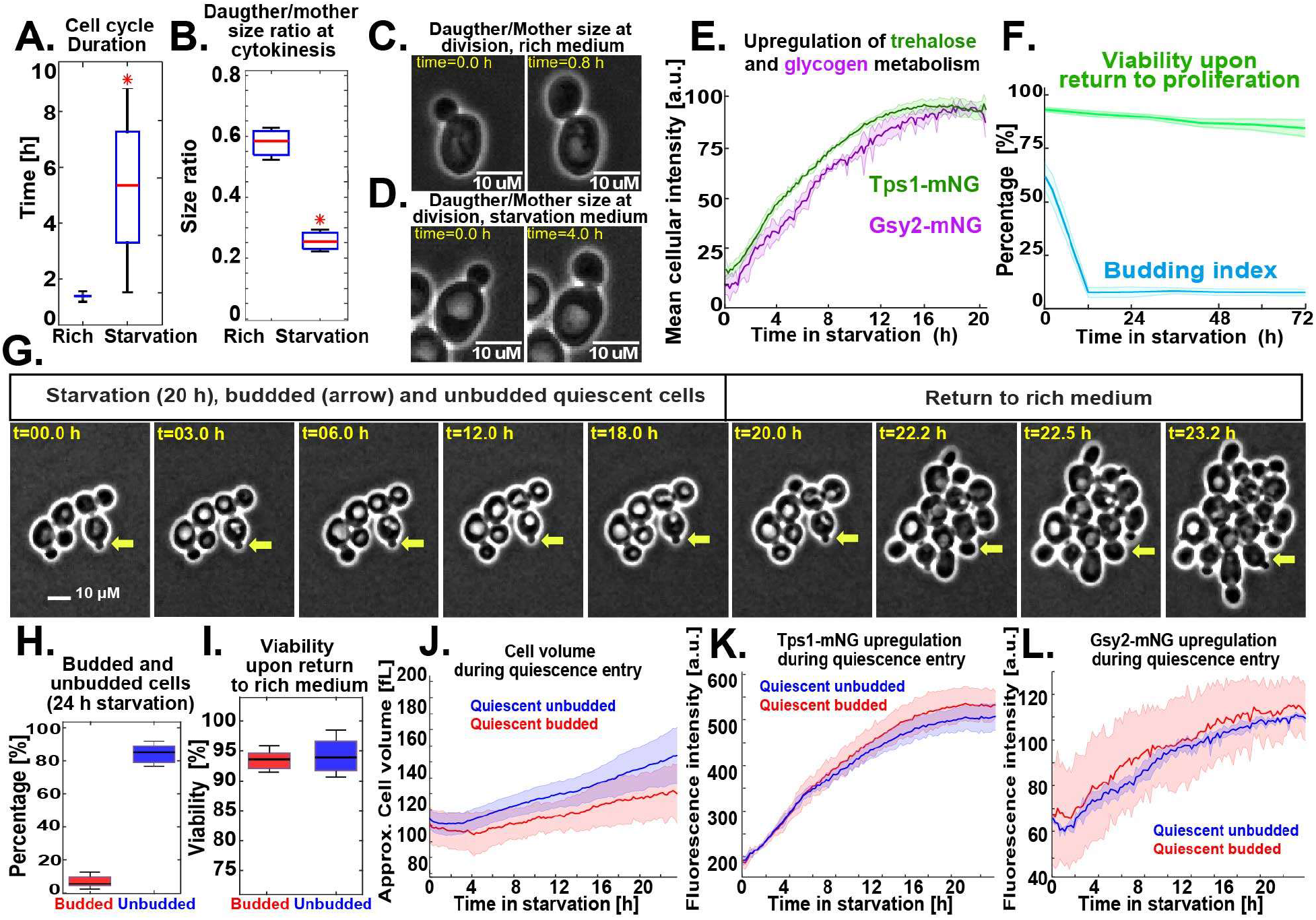
*S. cerevisiae* enters heterogeneous quiescent states under acute starvation. **(A)** Wild-Type cell cycle duration (*strain OAM128*) measured by the time between two consecutive cytokinesis events, in rich medium (n=13, 400 cells) or after exposure to acute starvation (n=9, 300 cells). **(B)** Daughter/mother cell size ratio at cytokinesis, measured as cell area in pixels, in rich medium (n=14, 400 cells) or for cells that completed cell cycle under starvation (n=9, 300 cells). (**C-D**) Representative phase contrast (PC) micrographs for cell division in **(C)** rich and **(D)** starvation medium. **(E)** Scaled mean cellular fluorescence intensity of the trehalose metabolism marker Tps1-mNeonGreen (n=9, 499 cells, *OAM394*), and the glycogen metabolism marker Gsy2-mNeonGreen (n=9, 300 cells, *OAM396*), during starvation. **(F)** Population budding index during starvation (blue) and viability (green) of quiescent cells upon return to proliferation (n=3, 237 cells). **(G)** Representative PC micrographs of cells during the starvation-proliferation transition showing a budded quiescent cell (yellow arrow). **(H-I)** Percentage of **(H)** budded and unbudded quiescent cells after 24 h starvation (n=6, 400 cells) and **(I)** their viability measured as proliferation resumption upon return to rich medium. **(J)** Average cell volume of quiescent budded and unbudded cells during starvation. **(K)** Average Tps1-mNG intensity in budded and unbudded quiescent cells. **(L)** Average Gsy2-mNG intensity in budded and unbudded quiescent cells. All data from biological replicates. Red star = p-value < 0.05, KS-test. Solid lines with shaded area = average plus 95% confidence interval. Boxplots: central mark, median; box bottom and top limit, 25 th and 75 th percentiles; whiskers, most extreme nonoutlier values.

### Acute starvation induces quiescence in states of low- or high-Cdk1 activity

To analyze the Cdk1 activity state of budded and unbudded quiescent cells, we tracked the formation of the septin ring at the bud site —which is absent during the low-Cdk1 state of G1 but present during the high-Cdk1 state of S-M phase— by C-terminally tagging the septin Cdc10 with the fluorophore mCyOFP1 (McMurray and Thorner, 2009). The presence of the septin ring was measured by the standard deviation of Cdc10 fluorescence at the cell periphery (Cdc10 Signal; Fig. 2 A, Fig. S1 B). To analyze the Cdc10 signal of single cells during the proliferation-quiescence transition, we developed the algorithm “Time series Profiling by Machine Learning” (TPML) which groups singlecell time series according to similarity of cell cycle arrest into hierarchically sorted heatmaps (Fig. 2 B, Fig. S1 C, D; Video 2). TPML analysis of Cdc10-mCyOFP1 time series established three major groups or “Cdc10-clusters” named according to whether cells reached a G1 cell cycle arrest during the proliferation-quiescence transition. Cells in cluster one — or “post-mitotic G1” — were in a high-Cdk1 state at starvation onset, as judged by the presence of the septin ring, and finished one last division before arresting in G1 (Fig. 2 C and D, blue). Cells in cluster two — or “Non-G1” — were also in a high-Cdk1 state at starvation onset but remained arrested in this state without reaching G1 conditions as judged by a persistent septin ring (Fig. 2 C and D, red). Cells in cluster three — or “direct G1” — were in the low-Cdk1 state of G1 at starvation onset and remained arrested in this state (Fig. 2 C and D, black).

**Fig. 2.**
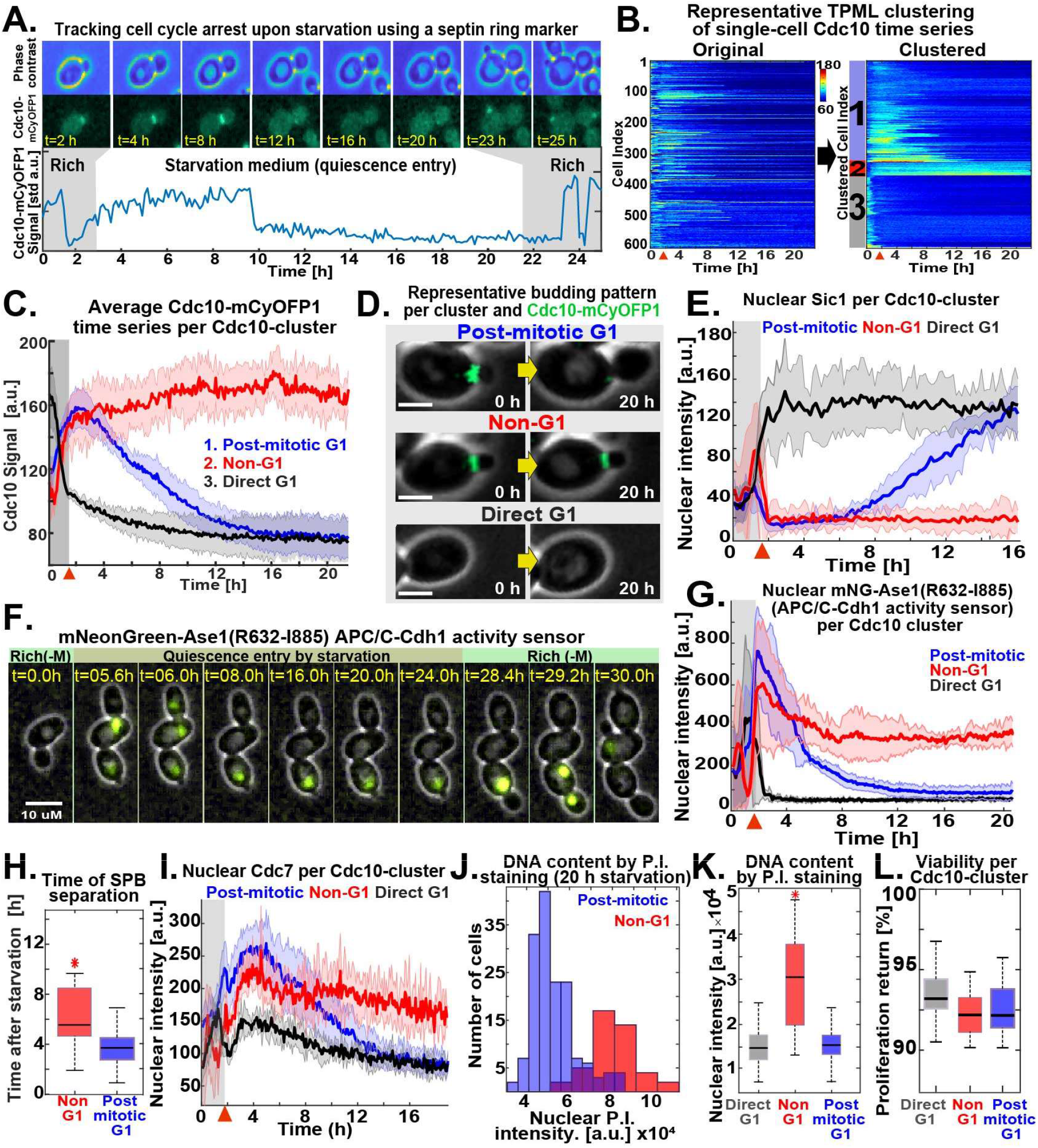
Acute starvation induces low- and high-Cdk1 quiescent states. **(A)** Schematic of cell cycle tracking during starvation using the septin Cdc10-mCyOFP1 signal (*OAM421*). **(B)** Representative heatmaps showing TPML arrangement of single-cell Cdc10 time series into Cdc10-clusters corresponding to the main patterns of cell cycle arrest upon starvation. **(C)** Average Cdc10 time series for each Cdc10-cluster (n=6, 633 cells). **(D)** Representative micrographs of budding patterns per Cdc10-cluster during starvation. Maximum intensity projection (MIP) mCyOFP1 image superimposed on phase contrast. Scale bar = 5 μM. **(E)** Average nuclear Sic1 intensity per Cdc10-cluster (n=3, 341 cells *OAM672*). **(F)** Representative micrographs of the APC/C-Cdh1 activity sensor mNeonGreen-Ase1(R632-I885) during starvation in post-mitotic (up) and Non-G1 (down) Q-cells (*OAM487*). MIP mNG image superimposed on phase contrast. R(-M) = Rich medium minus methionine. **(G)** Average mNG-Ase1(R632-I885) nuclear intensity per Cdc10-cluster (n = 3, 255 cells). **(H)** Time of spindle pole body (SPB) separation in Non-G1 and post-mitotic G1 Q-cells upon starvation (n = 3, 353 cells *OAM423*) **(I)** Average Cdc7-mScarlet-I nuclear intensity per Cdc10-cluster (n = 3, 345 cells *OAM422*) **(J)** Representative nuclear intensity histograms and **(K)** comparison of nuclear fluorescence intensity per Cdc10-cluster after propidium iodide (P.I.) staining (n=3, 304 cells *OAM421*). **(L)** Average viability of cells in each Cdc10-cluster upon return to rich medium after 24 h of starvation (n=3, 565 cells *OAM421*). All data from biological replicates. Red arrows = onset of starvation. Solid lines with shaded area = average ± 95% confidence intervals. Red star = p<0.05 by KS-Test. Boxplots display data from biological replicates: central mark, median; box bottom and top limit, 25 th and 75 th percentiles; whiskers, most extreme non-outlier values.

To assess the Cdk1 activity state of Cdc10-clusters, we imaged quiescence entry in cells bearing Cdc10-mCyOFP1 as a marker for the high-Cdk1 state of S-M phase, and mNG C-terminally tagged Sic1 as a marker for the low Cdk1-state of G1 (Hopkins et al., 2017). After starvation onset, nuclear Sic1 accumulated in post-mitotic G1 and direct G1 Q-cells according to their cell cycle progression (Fig. 2 E, black, blue). In contrast, nuclear Sic1 did not accumulate in Non-G1 Q-cells (Fig. 2 E, red), suggesting persistent Cdk1 activity. The high-Cdk1 kinase state of S-M phase is characterized by the inactivation of APC/C-Cdh1-mediated proteolysis (Zachariae et al., 1998). To measure the activity of APC/C-Cdh1 in Q-cells, we tracked the nuclear levels of the APC/C-Cdh1 activity sensor mNG-Ase1(R632-I885), expressed from the *MET3* promoter (Ondracka et al., 2016). During proliferation, the accumulation-degradation of the sensor followed the Cdc10 signal as expected for APC/C-Cdh1 substrates (Fig. S1 E). After starvation, the APC/C-Cdh1-sensor was degraded in post-mitotic G1 and direct G1 Q-cells (Fig. 2, F and G, black, blue) but persisted in Non-G1 Q-cells (Fig. 2, F and G, red), indicating APC/C-Cdh1 inactivation. Consistent with this observation, spindle pole body (SPB) separation, a process that requires APC/C-Cdh1 inactivation (Crasta et al., 2006), occurred, although with a delay, in Non-G1 Q-cells as judged by tracking of the SPB component Spc42 tagged with mTFP1 (Fig. 2 H).

To assess whether Non-G1 Q-cells entered S-phase, we tracked the DNA replication kinase Cdc7 C-terminally tagged with mScarlet-I (mSC). During starvation, nuclear Cdc7 persisted at higher levels in Non-G1 Q-cells (Fig. 2 I, red) in comparison to the other clusters (Fig. 2 I, blue, black). To measure DNA content in Q-cells, Cdc10-mCyOFP1 cells were microfluidically stained with propidium iodide after 20 hours in starvation medium. On average, Non-G1 Q-cells had duplicated DNA content (Fig. 2, J and K).

Together these results indicated that Non-G1 Q-cells persisted in a state of high-Cdk1 activity as judged by APC/C-Cdh1 inactivation, SPBs separation, DNA replication, and lack of Sic1 accumulation. High-Cdk1 quiescent states were equally viable to low-Cdk1 quiescence, as judged by comparing proliferation resumption among Cdc10-clusters (Fig. 2 L). We concluded that proliferating *S. cerevisiae* cells challenged by acute starvation after exiting G1 faced a decision-making process between remaining arrested in a high-Cdk1 state or completing a final cell cycle to reach a low-Cdk1 quiescent state in G1. In the rest of this study, we focused on cells experiencing starvation after G1 exit (postmitotic and Non-G1 Cdc10-clusters) and used the TPML algorithm to classify them into “low-Cdk1 quiescence” if they finished one last division or “high-Cdk1 quiescence” if they remained arrested.

### Low- and high-Cdk1 quiescent states share a core of stress responses but differ in the nuclear accumulation of Sfp1, Gln3, and Xbp1

To study how single cells challenged by starvation after G1 exit enter low- or high-Cdk1 quiescent states, we used a six-color fluorescent imaging system to assess to what extent differences in cell cycle progression or stress-activated factors determined high-Cdk1 quiescence entry (Arguello-Miranda et al., 2018). Imaging was optimized for using Cdc10-mCyOFP1 as cell cycle marker while simultaneously tracking five reporters for the major stress responses associated with the establishment of quiescence (Fig. 3 A, Fig. S2 A-H, Video 3). Metabolic stress was measured by the nuclear translocation of the nitrogen stress transcriptional regulator Gln3 (Crespo et al., 2002) and the carbon stress/retrograde pathway transcriptional regulator Rtg1 (Komeili et al., 2000). The biosynthetic capacity of the cell was tracked using the nuclear translocation of the ribosome biogenesis factor Sfp1 (Marion et al., 2004). Transcriptional repression was tracked using the histone deacetylase regulators Stb3 and Xbp1, which are key transcriptional repressors during quiescence (Mai and Breeden, 1997, McKnight et al., 2015).

**Fig. 3.**
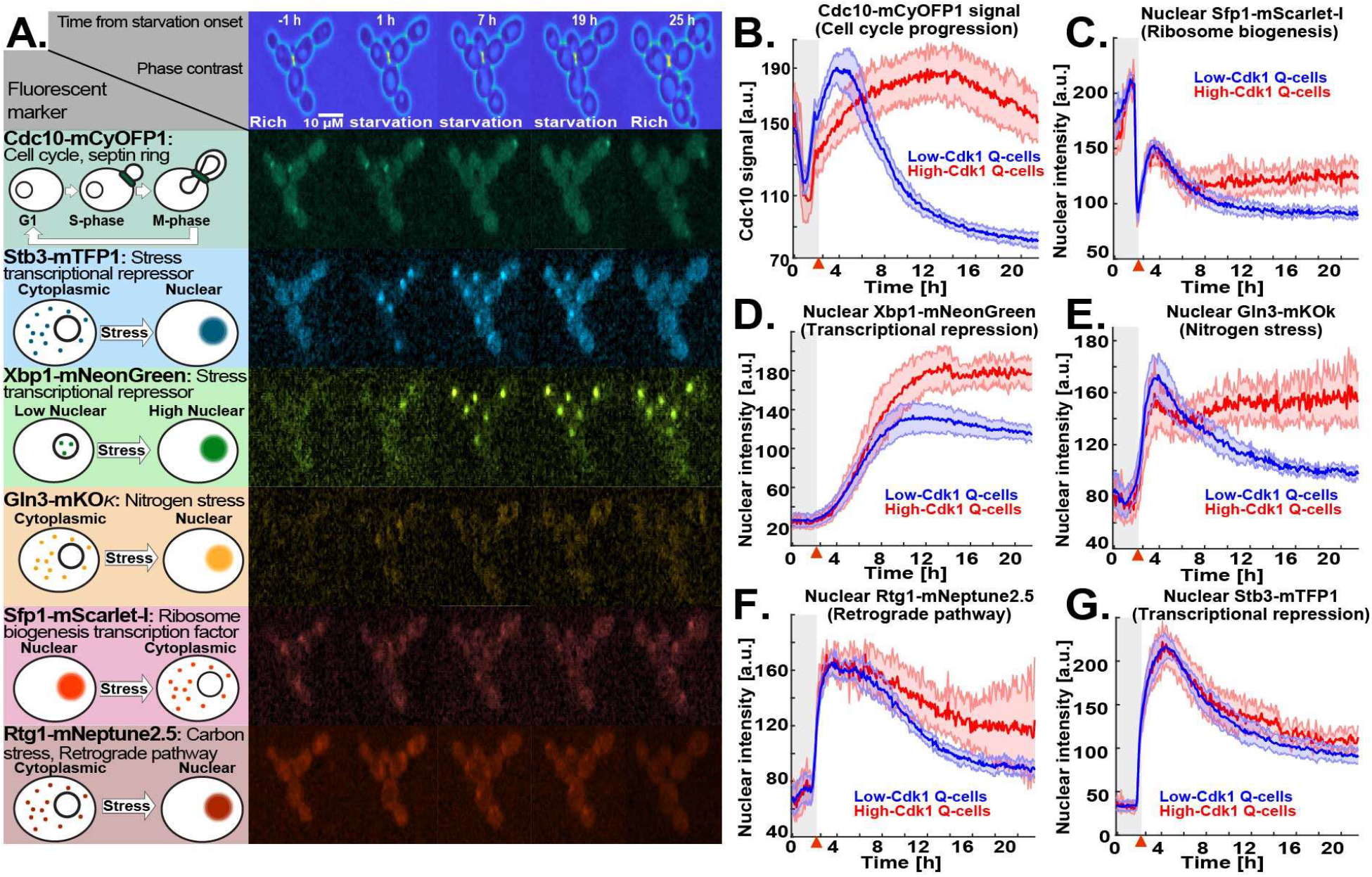
The stress transcription factors Sfp1, Gln3, and Xbp1 are selectively upregulated during high-Cdk1 quiescence entry. **(A)** Left, schematic nuclear accumulation of fluorescently tagged transcription factors used as sensors for stress responses. Right, representative MIP micrographs of the six fluorescent channels imaged in 6C1 cells upon starvation. **(B-G)** Quantification of cell cycle progression and stress responses in low-Cdk1 (blue) or high-Cdk1 (red) quiescent cells upon starvation (n=9, 825 cells *OAM425*). **(B)** Average Cdc10-mCyOFP1 time series per cluster **(C)** Average nuclear intensity of the ribosome biogenesis factor Sfp1-mSC per cluster. **(D)** Average nuclear intensity of the transcriptional repressor Xbp1-mNG per cluster. **(E)** Average nuclear intensity of the nitrogen stress response regulator Gln3-mKOk per cluster. **(F)** Average nuclear intensity of the carbon/retrograde pathway regulator Rtg1-mNeptune2.5 per cluster. **(G)** Average nuclear intensity of the transcriptional repressor Stb3-mTFP1 per cluster. Red arrows = onset of starvation. Solid lines with shaded area = average ± 95% confidence intervals.

Sfp1-Stb3-Rtg1-Gln3-Xbp1-Cdc10 (6C1) cells rapidly modulated the nuclear levels of stress markers upon starvation (Fig. S 2H). Low- and high-Cdk1 quiescent states were identified by TPML and the differences in the average time series of each group were assessed using the overlap in the 95% confidence interval. Strikingly, the most significant difference between low- and high-Cdk1 Q-cells was not cell cycle progression at starvation onset (Fig. 3 B), but the accumulation of nuclear Sfp1 (Fig. 3 C), Xbp1 (Fig. 3 D), and Gln3 (Fig. 3 E). In contrast, nuclear Rtg1 (Fig. 3 F) and Stb3 (Fig. 3 G) were accumulated to the same extent in low- and high-Cdk1 Q-cells.

To assess whether low- and high-Cdk1 Q-cells presented differences in stress responses not directly related to starvation, we imaged a second six-color (6C2) strain bearing fluorescent reporters for calcium signaling (Crz1-mTFP1) (Cyert, 2003), cell wall stress (Pkc1-mRuby3) (Mishra et al., 2017), osmoregulation (Hog1-mNeptune2.5) (Westfall et al., 2004), general stress (Msn2-mNG, Sfp1-mKOκ) (Granados et al., 2018), and cell cycle (Cdc10-mCyOFP1). 6C2 cells readily reported stress stimuli if perturbed during proliferation (Fig. S2 I). Upon starvation, the nuclear levels of Msn2 (Fig. 4 A), Crz1 (Fig. 4 B), Hog1 (Fig. 4 C), and the cell membrane-associated levels of Pkc1 (Fig. 4 D) were similar in low- and high-Cdk1 Q-cells.

**Fig. 4.**
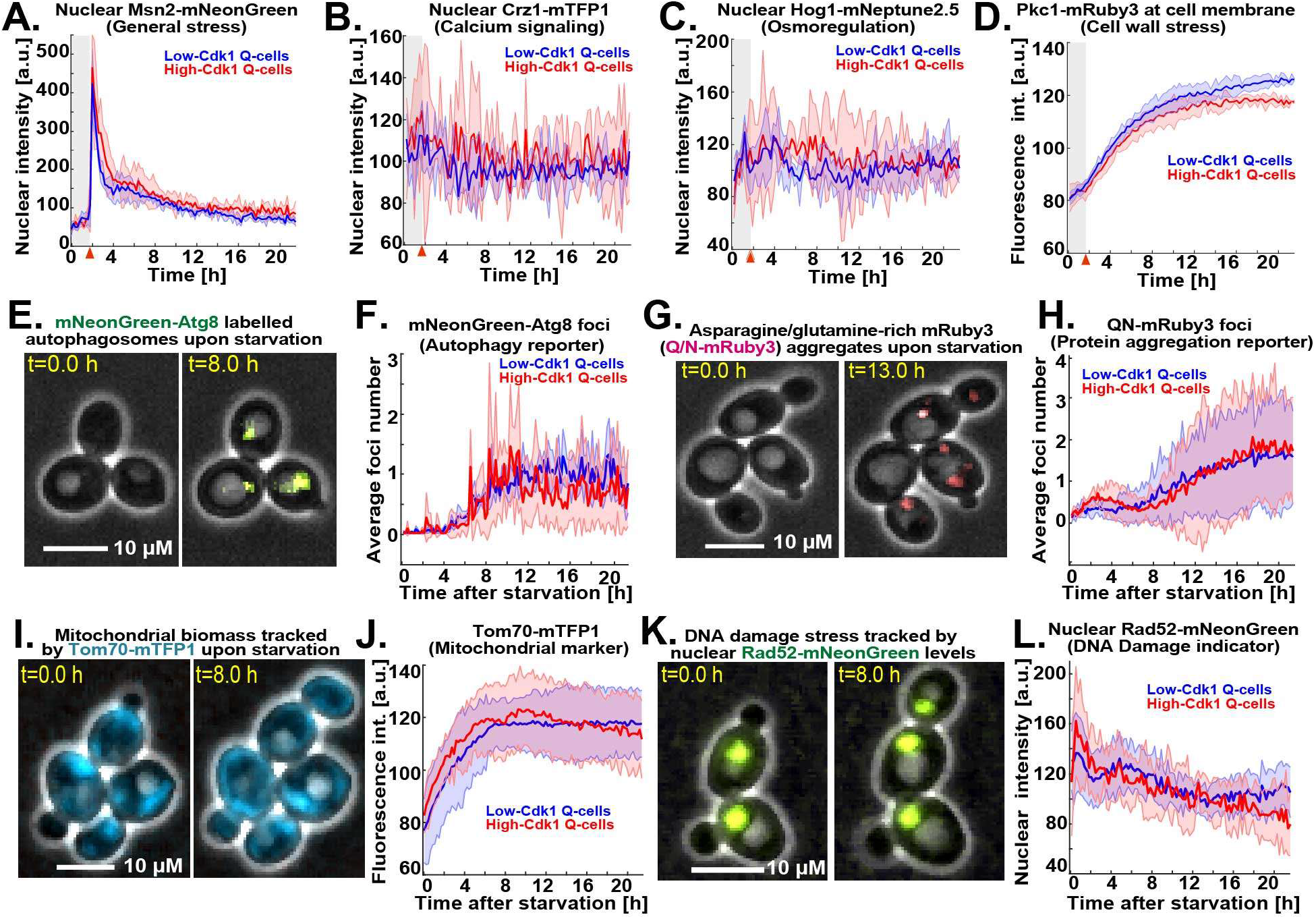
Low- and high-Cdk1 quiescent states share a core of stress-associated processes. **(A-D)** Quantification of non-metabolic stress responses in low-Cdk1 (blue) or high-Cdk1 (red) quiescent cells upon starvation (n=3, 255 cells *OAM667*). **(A)** Average nuclear intensity of the general stress transcription factor Msn2-mNG per cluster. **(B)** Average nuclear intensity of the calcium stress transcription factor Crz1-mTFP1 per cluster. **(C)** Average nuclear intensity of the osmotic stress kinase Hog1-mNeptune2.5 per cluster. **(D)** Average intensity of the cell wall stress kinase Pkc1-mRuby3 at cell membrane per cluster. **(E)** Representative micrographs of cells bearing the autophagy reporter mNG-Atg8 upon starvation. MIP mNG image superimposed on phase contrast. **(F)** Average number of mNG-Atg8 foci per cluster upon starvation (n=3, 450 cells *OAM654*). **(G)** Representative micrographs of cells bearing the protein aggregation sensor QN-mRuby3 upon starvation. MIP fluorescent mRuby3 image superimposed on phase contrast. **(H)** Average number of QN-mRuby3 foci per cluster upon starvation (n=3, 333 cells *OAM671*). **(I)** Representative micrographs of cells bearing the mitochondrial marker Tom70-mTFP1 upon starvation. MIP mTFP1 image superimposed on phase contrast. **(J)** Average total intensity of Tom70-mTFP1 per cluster upon starvation (n=3, 225 cells *OAM673*). **(K)** Representative micrographs of cells bearing the DNA damage marker Rad52-mNG upon starvation. MIP mNG image superimposed on phase contrast. **(L)** Average Rad52-mNG nuclear intensity per cluster upon starvation (n=3, 240 cells *OAM670*). Solid lines with shaded area = average ± 95% confidence intervals. Red arrows, onset of starvation.

To investigate the status of quiescence-associated processes (Sagot and Laporte, 2019) in low- and high-Cdk1 Q-cells, we induced starvation in Cdc10-mCyOFP1 strains containing fluorescent sensors for autophagy, protein aggregation, mitochondrial biomass, and DNA damage. Autophagy was measured by the foci number of an autophagosome marker, mNG-Atg8, expressed from its own promoter (Fig. S2 J) (Nair et al., 2011). Upon starvation, the number of Atg8 foci increased with no difference between quiescent states (Fig. 4, E and F). Protein aggregation was measured using a glutamine-asparagine-rich peptide (Antonets et al., 2016) bound to a mRuby3 fluorophore (QN-mRuby3) under the control of a copper-inducible promoter (*CUP1-1*). *CUP1p*-expression of QN-mRuby3 during proliferation produced few foci that remained within mother cells, as expected from protein aggregates in *S. cerevisiae* (Fig. S2 K) (Saarikangas et al., 2017). By contrast, expression of QN-mRuby3 during quiescence led to several foci per cell independent of cell cycle progression (Fig. 4, G and H; Fig. S2 M). Total mitochondrial biomass, as measured by the mean intensity of the translocase Tom70 (Hughes et al., 2016), showed no difference between quiescent states (Fig. 4, I and J). Rad52 nuclear levels and foci, which increase upon DNA damage (Alvaro et al., 2007), were not significantly different between quiescent states (Fig. 4, K and L; Fig. S2, L and N).

These results showed that although quiescent states shared a core of stress-associated processes, high-Cdk1 quiescence entry was characterized by increased nuclear accumulation of Sfp1, Gln3, and Xbp1.

### High-Cdk1 quiescence entry is determined by the stress status of the cell

The increased nuclear accumulation of Xbp1, Gln3, and Sfp1 in high-Cdk1 Q-cells suggested that the stress status of the cell, rather than differences in cell cycle progression, promoted high-Cdk1 quiescence entry upon starvation. To test this hypothesis, 6C1 cells were cell cycle-synchronized with a nocodazole arrest-release protocol before exposure to starvation (Fig. 5 A) and then clustered using TPML to identify cells with the same cell cycle synchrony around starvation onset (Fig. S3 A). This revealed that cells with identical synchrony, as judged by the Cdc10 signal overlap, could diverge into low- or high-Cdk1 quiescent states (Fig. 5 B; Fig. S3 B). A G1 synchronization protocol confirmed this result (Fig. S3 C). As in unsynchronized cultures, high-Cdk1 Q-cells accumulated higher nuclear levels of Xbp1 (Fig. 5 C), Gln3 (Fig. 5 D), and Sfp1 (Fig. 5 E), but not Rtg1 or Stb3 (Fig. S3 B). This showed that high-Cdk1 quiescence was not defined by cell cycle stage at starvation onset.

**Fig. 5.**
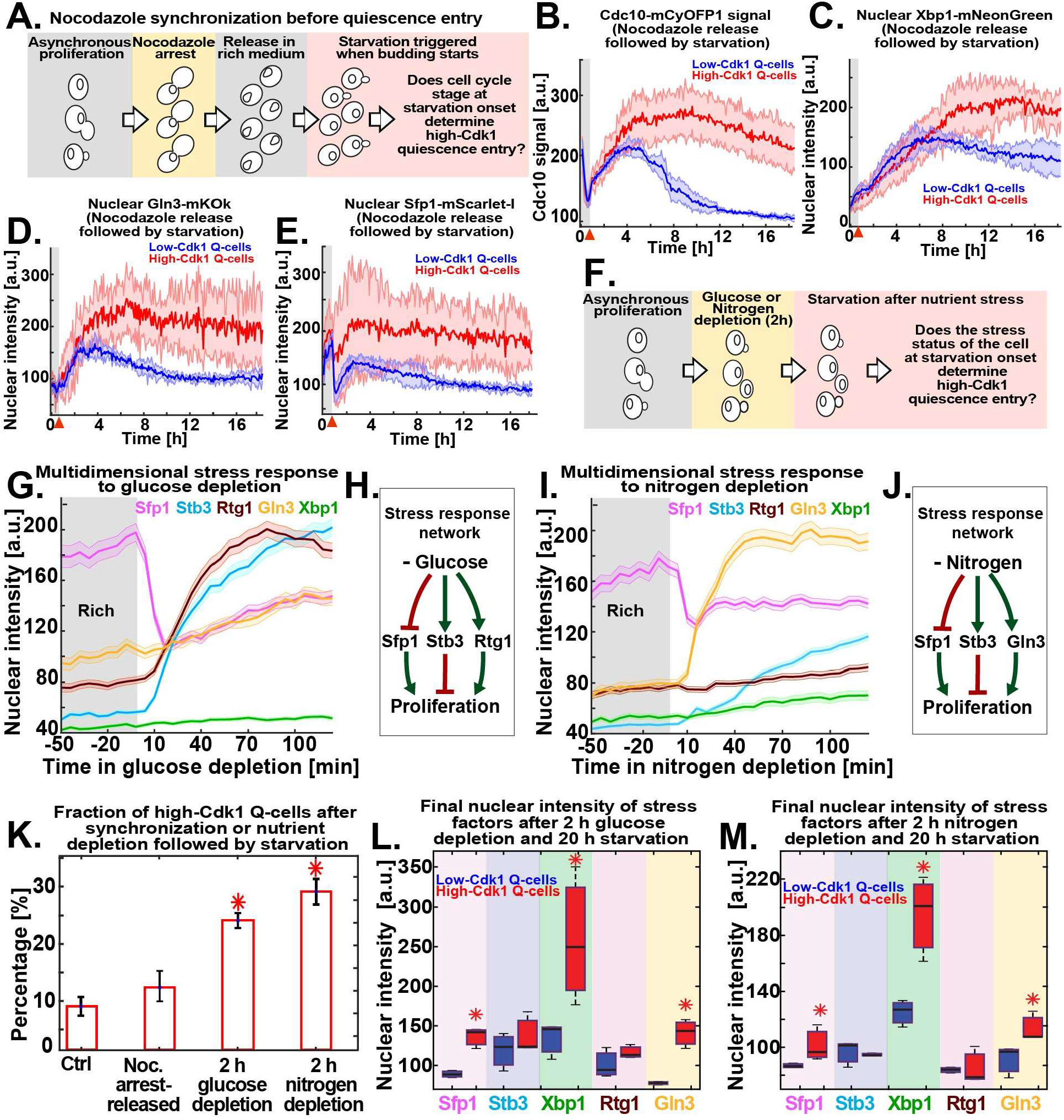
High-Cdk1 quiescence entry is determined by the stress status of the cell during starvation. **(A)** Schematic of nocodazole cell cycle synchronization before starvation onset. **(B-E)** Average time series for low-Cdk1 (Blue) and high-Cdk1 (red) Q-cells with the same cell cycle progression around starvation onset after nocodazole arrest-release. Average time series for **(B)** Cdc10 and the nuclear intensity of **(C)** Xbp1, **(D)** Gln3, and **(E)** Sfp1 in nocodazole arrest-released cells with same cell cycle progression at starvation onset (n=3, 96 cells). **(F)** Schematic of stress induction before starvation onset. **(G)** Nuclear intensity of stress factors as population average upon glucose depletion before starvation (504 cells). **(H)** Schematic of glucose-depletion induced stress response network. **(I)** Nuclear intensity of stress factors as population average upon nitrogen depletion before starvation (222 cells). **(J)** Schematic of nitrogen-depletion induced stress response network. **(K)** Percentage of high-Cdk1 Q-cells upon 20 h starvation in 6C1 cells with the following treatments before starvation: unsynchronized and unstressed control (Ctrl); nocodazole-synchronized (Noc.); 2 h glucose depletion; 2 h nitrogen depletion (n=7 per treatment) **(L-M)** Quantification of the final nuclear intensity of stress markers after 20 h starvation in low-Cdk1 (blue) and high-Cdk1 (red) Q-cells pre-treated with **(L)** glucose or **(M)** nitrogen depletion for 2 h before starvation (n=7, >233 cells). Solid lines with shaded area = average ± 95% confidence intervals. Red arrows = onset of starvation. Red star, p<0.05, KS-test. Bar plots = mean ± standard deviation. Boxplots display data from biological replicates: central mark, median; box bottom and top limit, 25 th and 75 th percentiles; whiskers, most extreme non-outlier values.

To assess whether high-Cdk1 quiescence was predisposed by the stress status of the cell at starvation onset, 6C1 cells were exposed to medium lacking glucose or nitrogen for two hours before starvation (Fig. 5 F). Glucose depletion generated a multidimensional stress response characterized by downregulation of nuclear Sfp1 and upregulation of Stb3 and Rtg1 (Fig. 5, G and H). Nitrogen depletion produced a different multidimensional stress response with nuclear translocations for Gln3, Sfp1, and Stb3 (Fig. 5, I and J). Triggering acute starvation from these two different stress states led to a significant increase of high-Cdk1 Q-cells (p<0.05, Fig. 5 K) with high final levels of nuclear Xbp1, Gln3, and Sfp1 (Fig. 5, L and M; Fig. S4, D and E).

Together, these experiments showed that high-Cdk1 quiescence entry was determined by the stress status of the cell and not by cell cycle stage at starvation onset.

### Nuclear levels of stress transcription factors predict high-Cdk1 quiescence

To assess the contribution of stress responses to the establishment of high-Cdk1 quiescence, we used linear discriminant classifiers (LDC) to find the best combinations of stress markers that discriminated between low- and high-Cdk1 quiescent states. To quantify the discrimination between quiescent states, the single-cell Mahalanobis distance to the mean of the low-Cdk1 Q-cells cluster was used. If a stress marker discriminates well between clusters, this metric should be small for low-Cdk1 Q-cells and large for high-Cdk1 Q-cells (Fig. 6 A). Time series obtained during quiescence entry in 6C1 (Fig. 3) and 6C2 (Fig. 4) cells were used as input for LDCs, and the average separation between clusters using single or combined parameters at each time point was calculated. LDCs based on Xbp1 and Gln3 could significantly discriminate between low- and high-Cdk1 quiescent states after ten hours into starvation (Fig. 6 B), whereas all other stress markers or their combinations, did not (Fig. 6 C; Fig. S3 F).

**Fig. 6.**
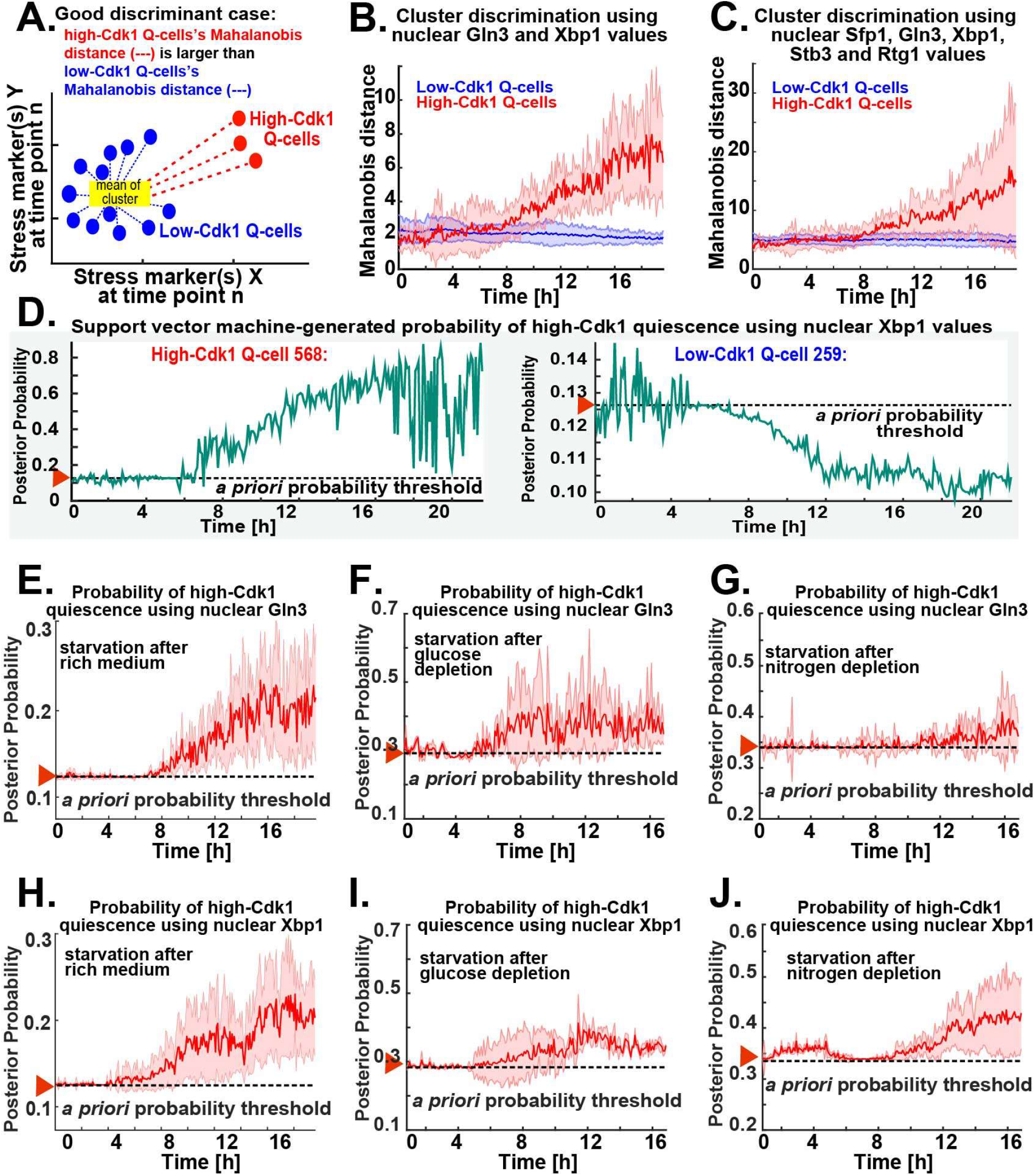
Nuclear accumulation of stress-activated transcription factors predicts high-Cdk1 quiescence entry depending on starvation conditions. **(A)** Schematic quantification of cluster discrimination by a linear discriminant classifier (LDC) based on the Mahalanobis distance. **(B-C)** Average cluster separation between low-Cdk1 (blue) and high-Cdk1 (red) Q-cells, as measured by the Mahalanobis distance, calculated using LDCs based on the combined nuclear intensity values of **(B)** Xbp1 and Gln3 or **(C)** all stress markers. **(D)** Probabilities of high-Cdk1 quiescence during starvation in a representative high-Cdk1 (left) and low-Cdk1 (right) Q-cell as measured by a Support Vector Machine Classifier (SVM) approach. **(E-G)** Average probability of high-Cdk1 quiescence assigned to high-Cdk1 Q-cells by SVMs based on the final nuclear intensity of Gln3 after exposure to **(E)** rich medium, **(F)** glucose depletion, or **(G)** nitrogen depletion before starvation. **(H-J)** Average probability of high-Cdk1 quiescence assigned to high-Cdk1 Q-cells by SVMs based on the final nuclear Xbp1 intensity after exposure to **(H)** rich medium, **(I)** glucose depletion, or **(J)** nitrogen depletion before starvation. Red arrow with horizontal dotted line, *a priori* probability threshold corresponding to the fraction of high-Cdk1 Q-cells in each experiment. Solid lines with shaded area = average ± 95% confidence intervals based on biological replicates (n>3).

To evaluate the predictive power of Xbp1 and Gln3, support vector machine classifiers (SVM) were used to calculate probabilities of high-Cdk1 quiescence using data from assays in which the 6C1 strain was exposed to rich medium (Fig. 3), glucose depletion (Fig. S3 D) or nitrogen depletion (Fig. S3 E) before starvation. SVMs were trained using the final nuclear values of Xbp1 or Gln3, and single-cell probabilities of high-Cdk1 quiescence upon starvation were computed for all time points (Fig. 6 D). Factors were considered predictive when correctly assigning probabilities of high-Cdk1 quiescence over the *a priori* probability threshold given by the fraction of high-Cdk1 Q-cells in the experiment.

Nuclear Gln3 produced predictions for high-Cdk1 quiescence over the *a priori* threshold only when cells were transferred from rich medium into starvation (Fig. 6 E) but not when cells were pre-treated with glucose (Fig. 6 F) or nitrogen depletion (Fig. 6 G) before starvation. In contrast, nuclear Xbp1 produced predictions for high-Cdk1 quiescence over the *a priori* threshold regardless of quiescence entry conditions (Fig. 6, H and I) and as early as 1 h into starvation for cells pre-treated with nitrogen depletion (Fig. 6 J). The average predictive power of Xbp1 across experiments, however, was significant after 8-9 h into starvation, suggesting that Xbp1 was correlated to high-Cdk1 quiescence when accumulated to high levels.

These results indicated that although different stress responses could contribute to high-Cdk1 quiescence depending on starvation conditions, increased nuclear Xbp1 was a general and potentially causative factor of high-Cdk1 quiescence.

### Nuclear accumulation of the transcriptional repressor Xbp1 promotes high-Cdk1 quiescence entry during starvation

To evaluate whether increased nuclear Xbp1 levels were causative of high-Cdk1 quiescence, while simultaneously measuring the stress status of the cell, we tracked quiescence entry in a Cdc10-mCyOFP1, Gln3-mKOκ, Rtg1-mNeptune2.5, Sfp1-mSC strain endogenously expressing Xbp1-mNG and also expressing Xbp1-mTFP1 from a *CUP1* promoter that is mildly induced by starvation (Fig. 7 A)(Peng et al., 2015). As an internal control for copper induction experiments, cultures were spiked with isogenic cells lacking *CUP1*p-*XBP1-mTFP1*, and both populations were algorithmically sorted using the mTFP1 signal before TPML analysis (Fig. S4 A).

**Fig. 7.**
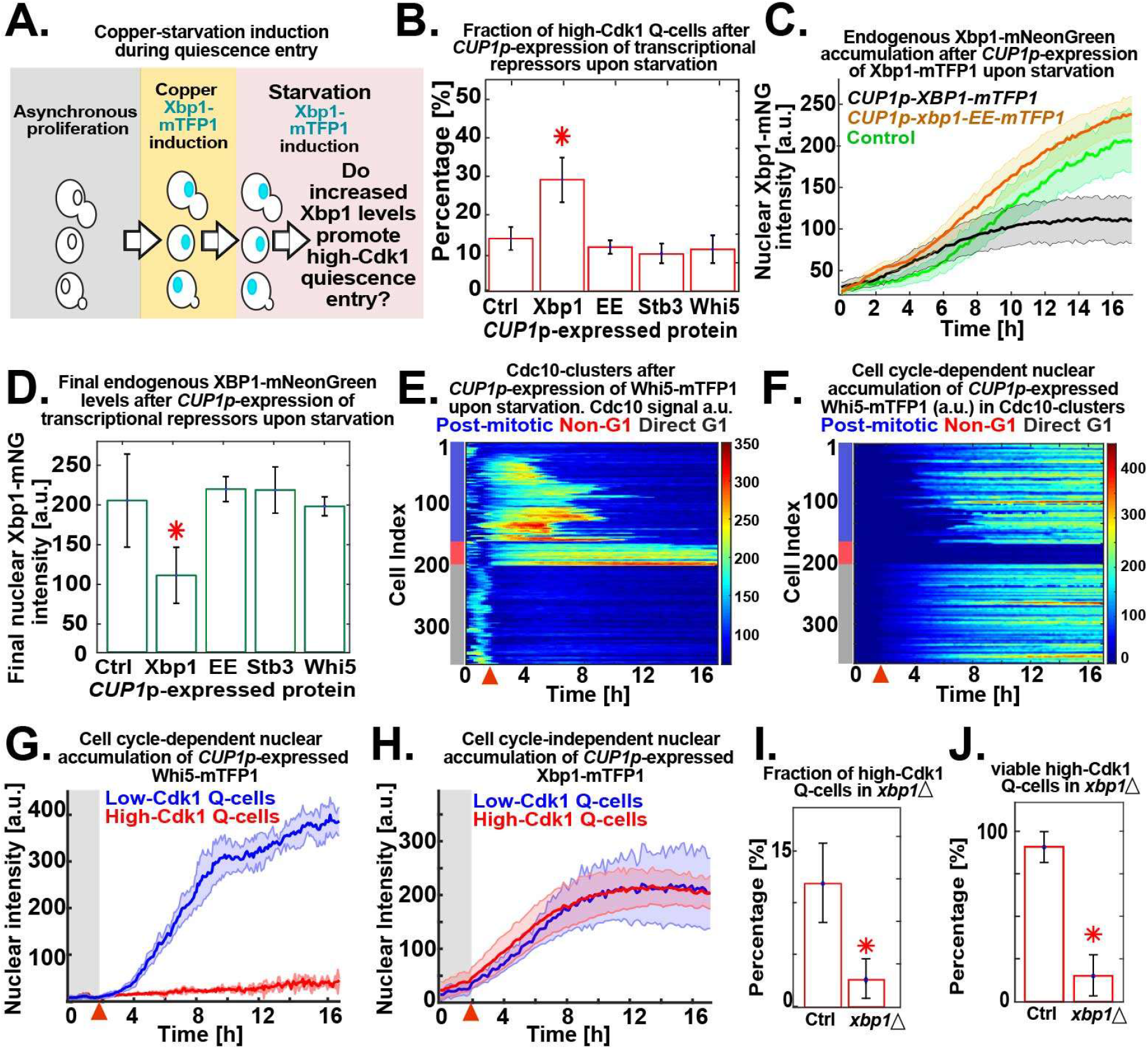
Nuclear accumulation of the transcriptional repressor Xbp1 promotes high-Cdk1 quiescence. **(A)** Schematic copper/starvation-induction protocol to assess the effect of increased Xbp1 levels during starvation. **(B)** Percentage of high-Cdk1 Q-cells after *CUP1p*-expression of mTFP1 C-terminally tagged wild-type Xbp1 (n=6, *OAM458*), the DNA binding domain mutant xbp1-EE (n=7, *OAM466*), the Xbp1-related transcriptional repressor Stb3 (n=4, *OAM497*), or the cell cycle repressor Whi5 (n=7, *OAM455*) during starvation. Ctrl = isogenic control strain (*OAM454*). **(C)** Nuclear accumulation of endogenously expressed Xbp1-mNG in cells expressing mTFP1-C-terminally tagged wild-type Xbp1 (n=6) or the DNA binding domain mutant xbp1-EE (n=7). **(D)** Final nuclear intensity of endogenously expressed Xbp1-mNG in cells expressing mTFP1 C-terminally tagged Xbp1 (n=6), xbp1-EE (n=7), Stb3 (n=4), or Whi5 (n=7) from the *CUP1* promoter during starvation. **(E-F)** Single-cell time series heatmaps contrasting the opposing pattern of **(E)** Cdc10-mCyOFP1 and **(F)** nuclear Whi5-mTFP1 after *CUP1p*-expression of Whi5-mTFP1 during starvation. **(G)** Average nuclear intensity of *CUP1p*-expressed Whi5-mTFP1 in low-Cdk1 (blue) and high-Cdk1 (red) Q-cells (n=7) **(H)** Average nuclear intensity of *CUP1p*-expressed Xbp1-mTFP1 in low-Cdk1 (blue) and high-Cdk1 (red) Q-cells (n=4) **(I-J)** Percentage of **(I)** high-Cdk1 Q-cells and **(J)** their viability, as measured by proliferation upon return to rich medium after 20 h of starvation, in *xbp1*Δ (n=4, 500 cells *OAM500*) and control *XBP1* cells (Ctrl, n=4, 311 cells *OAM502*). Red arrow = onset of starvation. Solid lines with shaded area = average ± 95% confidence intervals. Red star, p<0.05, KS-test. Bar plots = mean ± standard deviation.

*CUP1*p-expression of Xbp1-mTFP1 during starvation (Video 4) significantly increased the fraction of high-Cdk1 Q-cells (p<0.05, Fig. 7 B) without altering stress responses (Fig. S4 B). Interestingly, *CUP1*p-expressed Xbp1-mTFP1 reduced the levels of endogenously expressed Xbp1-mNG (Fig. 7, C and D), suggesting autoinhibition.

To test whether the effect of *CUP1p-XBP1-mTFP1* required its transcriptional repressor activity, we generated a DNA binding domain mutant of *XBP1* by introducing the mutations R349E and Q356E, which prevent DNA binding in related transcription factors (Mai and Breeden, 1997, Liu et al., 2015, Nair et al., 2003). *CUP1*p-expression of xbp1-EE-mTFP1 during starvation had no impact on the fraction of high-Cdk1 Q-cells (Fig. 7 B), the endogenous Xbp1-mNG levels (Fig. 7, C and D), or stress responses (Fig. S4 C). To determine the specificity of Xbp1’s effect in promoting high-Cdk1 quiescence, we used the *CUP1* promoter to express Stb3, which is a transcriptional repressor functionally related to Xbp1 (Liko et al., 2010), and Whi5, a major transcriptional repressor of cell cycle (Schmoller et al., 2015). *CUP1*p-expression of Stb3-mTFP1 or Whi5-mTFP1 did not alter the fraction of high-Cdk1 Q-cells (Fig. 7 B) or endogenous Xbp1 levels (Fig. 7 D) and had negligible effects on stress responses (Fig. S4, D and E).

The nuclear accumulation of *CUP1*p-expressed Whi5-mTFP1 strictly depended on cell cycle progression, as indicated by the opposing patterns of Cdc10 (Fig. 7 E) and Whi5 (Fig. 7 F) in Cdc10-clusters, remaining excluded from the nucleus of high-Cdk1 Q-cells (Fig. 7 G, Video 5). In stark contrast, nuclear accumulation of *CUP1*p-expressed Xbp1-mTFP1 was entirely independent of Cdk1 activity state (Fig. 7 H) or cell cycle progression (Fig. S4 B).

TPML analysis of *xbp1Δ* cells bearing markers for stress (Msn2-mNG, Gln3-mKOκ, Rtg1-mNeptune2.5, Sfp1-mSC-I, Stb2-mTFP1) and cell cycle progression (Cdc10-mCyOFP1) showed that deletion of *XBP1* led to altered patterns of cell cycle arrest during starvation (Video 6), reducing the total fraction (Fig. 7 I) and viability (Fig. 7 J) of high-Cdk1 quiescent Q-cells, without affecting the onset of stress responses (Fig. S7 F).

We concluded that Xbp1 was a causative factor of high-Cdk1 quiescence entry, which was sufficient to increase the frequency of high-Cdk1 Q-cells when accumulated at high levels.

### Nuclear Xbp1 acts as a time-delayed integrator to record single-cell history of stress stimuli

The capacity of Xbp1 to promote quiescent states suggested that it must be rapidly downregulated upon return to proliferation. To test this hypothesis, we analyzed the nuclear levels of stress transcription factors in 6C1 cells that were returned to rich medium after 20 h of starvation. Surprisingly, we found that whereas stress factors such as Sfp1 or Stb3 readily translocated upon exposure to rich medium, the nuclear levels of Xbp1 remained invariable for 2 h ± 20 min after exposure to rich medium (Fig. 8 A; Fig. S5 A) and after that decreased by dilution through cell division (Fig. 8 B; Fig. S5 B). This indicated that Xbp1 was slowly downregulated after return to rich medium, suggesting that information from past quiescent states persisted in the nuclear levels of Xbp1.

**Fig. 8.**
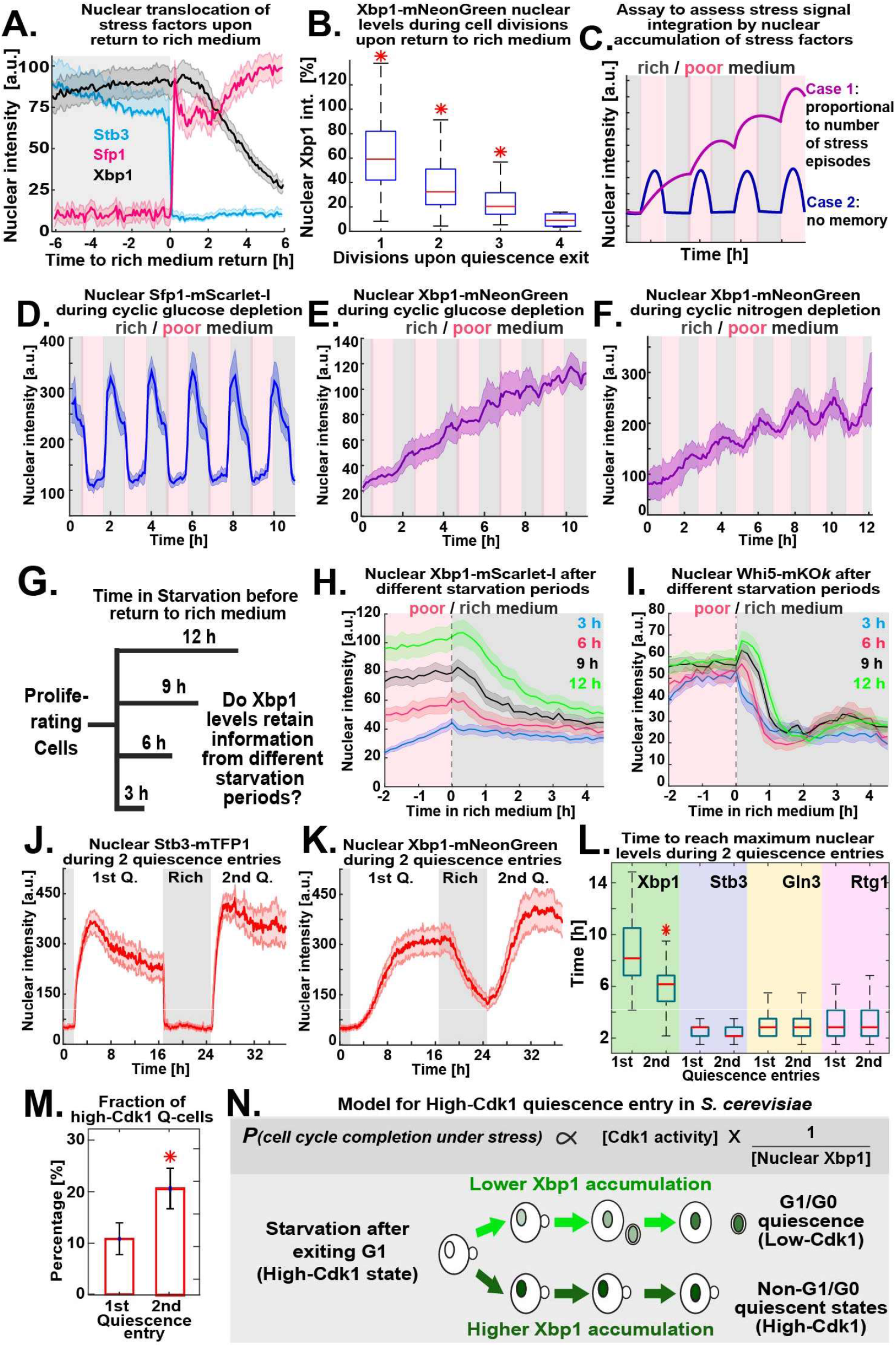
Xbp1 acts as a time-delayed integrator of stress stimuli. **(A)** Scaled average nuclear intensity of Sfp1-mScarlet-I, Stb3-mTFP1 and Xbp1-mNG during return to proliferation after 20h in starvation (n=6, 210 cells *OAM425*) **(B)** Boxplots of nuclear Xbp1 intensity, as percentage of nuclear Xbp1 intensity at 20 h starvation, during cell divisions after return to proliferation (n=6, > 101 cells *OAM425*). **(C)** Schematic assay to assess stress stimuli integration by nuclear accumulation of stress factors. **(D-E)** Average nuclear **(D)** Sfp1 and **(E)** Xbp1 intensity during hourly episodes of glucose depletion (n=5, 335 cells). **(F)** Average nuclear Xbp1 intensity during hourly episodes of nitrogen depletion (n=5, 235 cells *OAM425*). **(G)** Schematic assay to assess the integration of starvation duration by nuclear accumulation of stress factors. **(H)** Nuclear Xbp1-mScarlet-I average intensity in cells transferred to rich medium after different starvation periods (n=3, >89 cells *OAM849*) **(I)** Nuclear Whi5-mKOk average intensity in cells transferred to rich medium after different starvation periods (n=3, >64 cells *OAM850*) **(J-K)** Average nuclear intensity of **(J)** Stb3 and **(K)** Xbp1 during two consecutive quiescence entries (n=4, 310 cells *OAM425*). **(L)** Boxplots of time to reach maximum nuclear intensity by stress factors during two quiescence entries (n=4, 310 cells *OAM425*). **(M)** Percentage of high-Cdk1 Q-cells during two consecutive quiescence entries (n=4, 310 cells *OAM425*). **(N)** Schematic model of high-Cdk1 quiescence entry. Top: probability of cell cycle completion under starvation (*P*) is proportional to Cdk1 activity (given by cell cycle stage at starvation onset) and inversely proportional to nuclear Xbp1 levels (given by single-cell history). Bottom: depending on Xbp1 accumulation, cells with identical cell cycle progression at starvation onset enter different quiescence states. Solid lines with shaded area = average ± 95% confidence intervals. Red star p<0.05, KS-test. Bar plots = mean ± standard deviation. Boxplots display data from biological replicates: central mark, median; box bottom and top limit, 25 th and 75 th percentiles; whiskers, most extreme non-outlier values.

To test whether information from past stress stimuli was recorded in the nuclear levels of Xbp1, we exposed 6C1 cells to hourly changes between rich medium (SCD) and either glucose-depleted medium or nitrogen-depleted medium (Fig. 8 C). Under these conditions, stress responses oscillated (Fig. 8 D; Fig. S5, C and D; Video 7), but only the nuclear levels of Xbp1 increased in proportion to the number of previous stress episodes (Fig. 8 E and F; Fig. S5, C and D).

To test whether nuclear Xbp1 levels retained information from past starvation episodes, we tracked Xbp1 in cells exposed to starvation for 3, 6, 9, or 12 h before return to rich medium (Fig. 8 G). Starvation periods of increasing duration produced populations that maintained significantly different Xbp1 levels after transfer to rich medium, with the 3 h and 12 h starvation-exposed populations maintaining different Xbp1 levels beyond 4 h after return to rich medium (*p*<0.05, Fig. 8 H, cells aligned to the time of return to rich medium). This behavior was analogous to a time-delayed integrator in control theory (the output signal —nuclear Xbp1 levels— corresponds to the time integral, or duration in this case, of the input signal —starvation period— (Hao et al., 2013)). To test whether other histone deacetylase regulators could act as temporal integrators of starvation signals, we tracked nuclear Whi5 and Stb3 levels in cells exposed to different starvation periods before return to rich medium; however, Whi5 (Fig. 8 I) and Stb3 (Fig. S5 E) nuclear levels did not reflect starvation duration.

A time-delayed integrator role for Xbp1 predicted that previous quiescence entries could increase the frequency of high-Cdk1 Q-cells in future quiescence entries by producing populations with higher Xbp1 levels. To test this hypothesis, we imaged 6C1 cells during two quiescence entries separated by a period in rich medium (Video 8). The nuclear translocation of stress transcription factors, such as Stb3 (Fig. 8 J), was not affected by a previous quiescence entry. In contrast, nuclear Xbp1 persisted in rich medium after the first quiescence exit (Fig. 8 K) and reached maximum accumulation 2 h ± 29 min faster during the second quiescence entry (Fig. 8 L). TPML Cdc10-clustering according to the first or second quiescence entry (Fig. S5 F) showed that the fraction of high-Cdk1 Q-cells significantly increased during the second quiescence entry (Fig. 8 M). This effect was abolished if the experiment was performed in *xbp1Δ* cells (Fig. S5 G).

These results showed that Xbp1 acted as an integrator of previous stress stimuli, which rendered the propensity of high-Cdk1 quiescence dependent on single-cell history.

## Discussion

In this study, we showed how proliferating *S. cerevisiae* populations challenged by acute starvation enter quiescent states with low- or high-Cdk1 kinase activity. The halt of proliferation upon acute starvation was accompanied by increased storage carbohydrate metabolism, mitochondrial biomass, protein aggregation, autophagy, stress resistance, and size asymmetry during cell division. This indicates that the general processes of quiescence entry observed in batch cultures can also be triggered by starvation under constant environmental conditions.

The use of machine learning classifiers allowed us to *in silico* isolate and characterize cells entering quiescence in a high-Cdk1 state. By turning the values of single or combined stress markers into probabilities of cell fate, this approach distinguished correlated (Sfp1 and Gln3) from causative (Xbp1) factors of high-Cdk1 quiescence. Our results show that stress responses are remarkably homogenous during the initial stages of quiescence entry and that cells can diverge into different quiescent states after starvation stimuli are integrated by the histone deacetylase regulator Xbp1. Although Xbp1 represses crucial cell cycle genes, such as cyclins *(CLN3)* and mitotic transcriptional activators *(NDD1)* (Miles et al., 2013), it has been considered dispensable for quiescence because *xbp1Δ* cells still access G1/G0 quiescence in batch cultures (Mai and Breeden, 1997); however, we find that Xbp1 is essential for viable high-Cdk1 quiescence.

The role of Xbp1 as a cell cycle-independent integrator of past stress signals shows that quiescence entry in *S. cerevisiae* depends on single-cell history. Remarkably, this implies that each cell can have a different point of commitment to finish cell cycle under stress, or restriction point (Johnson and Skotheim, 2013, Zetterberg et al., 1995), determined not only by Cdk1 activity (given by cell cycle stage at starvation onset) but also by nuclear Xbp1 levels (given by single-cell history) (Fig8. N).

The mechanism to control Xbp1 activity is unknown but likely involves auto-inhibition when accumulated to high levels (Fig. 7 C). Interestingly, quiescence in hair follicles relies on an inhibitory network motif whereby a Cdk1 inhibitor, p21, transcriptionally represses another Cdk1 inhibitor, p15 (Lee et al., 2013). Auto-inhibition or mutual inhibition of quiescence-promoting factors might be a universal mechanism to ensure the reversibility of the quiescent state.

How can cells persist in high-Cdk1 quiescence for a long time? Due to the bistable nature of cell cycle networks, the low-Cdk1 state of G1 and the high-Cdk1 state of S-M phase are stable-steady states, which can be, by definition, sustained for prolonged periods (Novak et al., 2007). We envision that, under persistent starvation, high levels of Xbp1 force cells to settle in the available stable-steady state by inhibiting proliferation-promoting positive feedback loops through transcriptional repression. Consistent with this, high-Cdk1 quiescence is promoted by high levels of Xbp1 and abolished by *XBP1* deletion. Increased nuclear Xbp1, however, did not stop *all* cells from completing cell cycle under acute starvation (Fig. 7B), indicating that other events, such as completion of DNA replication under stress or accumulation of mitotic cyclins, also regulate high-Cdk1 quiescence entry.

The control of quiescence entry by cell cycle-independent integrators of stress stimuli could provide an adaptive advantage in environments with strong fluctuations of nutrients, as is the case for parasites with complex life cycles and cancer cells attempting metastasis (Wang et al., 2015). In mammalian cells, transcriptional repressors that could play an analogous role to Xbp1 are HES1 (Sueda et al., 2019, Coller, 2011), the DREAM complex (Bainor et al., 2018, Miles and Breeden, 2017, Schade et al., 2019), and developmentally regulated histone deacetylase-regulators (Wang et al., 2019); however, it is unknown whether they integrate stress stimuli during quiescence entry.

In this study, we have characterized, using microfluidics and machine learning algorithms, the emergence of low- and high-Cdk1 quiescent states in proliferating S. *cerevisiae* cells challenged by acute starvation. Our results outline the temporal integration of stress stimuli through cell cycle independent transcriptional repressors as a mechanism to establish cellular quiescence outside of G1/G0.

## Materials and Methods

### Media

Rich medium was SCD (1% succinic acid, 0.6% sodium hydroxide, 0.5% ammonium sulfate, 0.17% yeast nitrogen base without amino acids or ammonium sulfate, 0.13% amino acid dropout powder (complete), 2% glucose). Glucose-depleted medium was SCG (SC with 2% glycerol instead of glucose). Nitrogen-depleted medium was SCD minus ammonium sulfate and amino acid dropout powder. Starvation medium was 0.6% potassium acetate, delivered at 0.6 psi flow rate, pH 7.2 adjusted with 0.125 M Na_2_CO_3_. For copper expression of proteins, cells were maintained in SCD plus 50 μM bathocuproine disulfonate (BCS) (Gross et al., 2000) until before loading in the microfluidic device. *CUP1p*-expression was triggered with SCD containing 25 μM CuSO_4_, added as a 1: 1000 dilution from a 25 mM stock solution (Labbe and Thiele, 1999). *MET3p*-expression was triggered with SCD lacking methionine. 200 mM CaCl_2_-containing SCD had NH_4_SO_4_ replaced by NHCl_3_.

### Plasmid Construction

All plasmids were confirmed by sequencing and restriction analysis. Table S1 lists the used primers and Table S2 lists the used plasmids. The *MET3*p-mNeonGreen-Ase1(R632-I885) plasmid resulted from a quadruple ligation of the following fragments: (1) a *Sac*I-*Pac*I flanked *MET3* promoter sequence (493-1 to -1) (2) a *Pac*I-*Asc*I flanked mNeonGreen (Yeast Optimized) fluorophore released from pYLB10 (Arguello-Miranda et al., 2018) (3) an *Asc*I-*Not*I flanked C-terminus of *ASE1* (R632-I885) 4) a pRS305 backbone cut with *Not*I and *Sac*I. The *MET3* promoter and *ASE1* fragment were amplified from WT W303 genomic DNA. Linearization with Age*I* allowed Integration at *LEU2*. pRS304 carrying *ATG8p-mNeonGreen-ATG8-ATG8ter* was created by replacing the *Pac*I-*Bam*HI flanked yeGFP sequence in *pRS304-ATG8p-2xyeGFP-ATG8-ATG8ter* (a gift from the Henne Lab, University of Texas Southwestern Medical Center) by a *Pac*I-*Bam*HI flanked mNeonGreen fluorophore amplified from pYLB10 (Arguello-Miranda et al., 2018). The pRS306 plasmid carrying *CUP1p-QN-mRUBY3* was created by triple ligation of the following fragments: (1) a *Sal*I-*Hind*III flanked *CUP1-1* promoter sequence (434-16 bp upstream of start codon) (2) a *Hind*III-*Pac*I flanked Q/N-rich sequence of *GLN3* (aa166-242) (3) a *Pac*I-*Sal*I pLondon266 backbone containing the mRuby3 fluorophore. *CUP1p* and *GLN3* (aa166-242) fragments were amplified from W303 genomic DNA. Linearization with *Stu*I allowed integration at *URA3. CUP1* expression plasmids were created by replacing the *Pac*I-*Asc*I flanked mCherry fluorophore in pLondon266 (Arguello-Miranda et al., 2018) with a *Pac*I-*Asc*I flanked mTFP1 fluorophore. The resulting pLondon266-mTFP1 was cut with *Sal*I-*Pac*I and triple-ligated to a to a *Sal*I-*Hind*III flanked *CUP1* promoter and the *Hind*III-*Pac*I flanked ORF of *XBP1, STB3*, or *WHI5*, which were amplified from W303 genomic DNA. Linearization with *Stu*I allowed integration at the *URA3. CUP1p-xbp1-EE-mTFP1* was created by swapping the DNA binding domain of *XBP1*, which is naturally flanked by *Cla*I-*Bsa*BI sites in S. *cerevisiae* W303, by a *Cla*I-*Bsa*BI fragment with arginine 349 and glutamine 356 changed to glutamic acid by PCR mutagenesis with the appropriate primers.

### Strain Construction

Strains were diploid S. *cerevisiae* W303 *(leu2-3,112 his3-11,15 ura3-1 trp1-1 can1-100 ade2)* (Table S3). Diploids were created by mating appropriate haploids obtained by tetrad dissection or by transformation with PCR tagging cassettes, deletion cassettes, or plasmids using the PEG/lithium acetate protocol (Longtine et al., 1998). *xbp1Δ::KANMX4* deletion cassette was amplified from BY4741 genomic DNA. C-terminal PCR tagging cassettes were amplified from Longtine (pFA6a) plasmids (Table S2). Tagged strains were confirmed by DNA sequencing, PCR, or functional assays. Tagged fluorescent proteins were heterozygous unless otherwise stated.

### Microfluidic cell culture

A Y04C CellASIC microfluidic device (http://www.cellasic.com/) operated at 25 °C, 0.6 psi flow rate, was used for culturing cells in all experiments. 50 μL of sonicated (4-6 s at 3 W) liquid SCD culture, OD_600_ = 0.3, were loaded with 1-2 pulses of 8 psi for 5 s. All medium changes were isobaric. To trigger quiescence, cells grew for 2 or 3 h in SCD before exposure to starvation medium for 20 h. For *MET3*p protein induction, cells grew 1 h in SCD before exposure to SCD minus methionine for 2 h. For *CUP1*p induction, cells were maintained in 50 μM of bathocuproine disulfonate (BCS)-containing SCD before exposure to 25 μM CuSO_4_-containing SCD for 2 h. For osmoregulatory tests, cells grew 2 h in SCD before exposure to 200 mM CaCl_2_-containing SCD for 30 min.

### Microfluidic staining of DNA

After 20 h in starvation, cells in the YO4C microfluidic device were treated at 25°C, 0.6 psi flow rate with: *(1)* a solution of 4M LiCl containing 50 mM Nystatin (from a stock solution of 3 mg/ml Nystatin in DMSO) for 4 h; *(2)* a solution of 50 mM Tris/HCl, pH 7.8, containing 0.3 mg/ml RNAse A (from a 10 mg/mL RNAse stock solution in 10mM sodium-acetate, pH 5.2) for 4 h; *(3)* FACS buffer (180mM Tris/HCl, 190mM NaCl, 70 mM MgCl2) containing 1.85 μM propidium iodide (added before use from a 0.5 mg/ml solution) for 2 h. Nuclear fluorescent intensity was measured in segmented cells as describe below.

### Statistical analysis

All statistical analyses were done in MATLAB and considered each lane of the microfluidic device as a biological replicate. Statistical significance was assessed using the Kolmogorov-Smirnov test using a 0.05 p-value. Bar plots display mean and standard deviation. Boxplots display the median as a central mark, the 25 th and 75 th percentiles are the bottom and top limit of the box, and the whiskers are the most extreme non-outlier values. Outliers were excluded using the function *isoutlier()*. Average times series were plotted as a solid line surrounded by a shaded area corresponding to 95 % confidence intervals defined by biological replicates.

### Time-Lapse Microscopy

A motorized, temperature-controlled, ZEN software-operated Zeiss Observer Z1 microscope with Definite Focus 2.0 was used. A 40X Zeiss EC Plan-Neofluar 40X 1.3 NA oil immersion objective and an AxioCam HR Rev 3 camera were used for image acquisition. At least five fields of view were imaged per biological replicate using a 6- or 12-min acquisition rate. A set of custom dichroic mirrors and bandpass filters were used for simultaneous imaging of six different fluorophores (Table S4) (Arguello-Miranda et al., 2018). Fluorophores and exposure times were: mCyOFP1 35 ms, mTFP1 400 ms, mNeonGreen 100 ms, mKOκ 400 ms, mScarlet-I 120 ms, mNeptune2.5 400 ms, and phase contrast 20 ms. *CUP1*p-expressed fluorescent proteins were imaged using 20 ms. LED light sources were kept at 12.5 % intensity (25% for mNeptune2.5). Correction for crosstalk between fluorescent channels was assessed using single-fluorophore tagged strains by calculating the cross-correlation between pixels in the signal-containing channel and the empty channels, the code for crosstalk detection is available at GitHub: https://github.com/alejandrolvido/Spectral-Imaging. Crosstalk corrections were: for strains OAM421 and OAM425, the corrected mKOκ image equals original mKOκ - 0.23 mCyOFP1, and for strains OAM455/458/466/497 the corrected mCyOFP1 image equals original mCyOFP1 - 0.33 mTFP1.

### Image processing and quantification of cellular features

The code for image analysis, single-cell quantifications, and time series visualization is available at https://github.com/alejandrolvido/Quiescence-Entry. The code is organized in subfolders corresponding to each main figure of this work. Images were collected using ZEN pro software (Zeiss) with 2 x 2 binning, exported in non-compressed TIFF format, and converted to double format before quantification. Shot noise was reduced using a *medfilt2()* filter with a 3 x 3 structuring element. After segmenting cells with published algorithms (Doncic et al., 2013, Wood and Doncic, 2019), image background was calculated as the median intensity of the space not occupied by cells. A 2D Gaussian fit to the brightest pixel of nuclear proteins was used to calculate nuclear parameters (Doncic et al., 2013), where nuclear intensity is defined as the mean intensity of nuclear pixels minus the mean intensity of cytoplasmic pixels. The septin ring was detected by measuring the standard deviation Cdc10-mCyOFP1 at cell periphery “Cdc10 Signal” (Figure S1B).

### Time series profiling by machine learning (TPML) algorithm

This algorithm aims to use a small number of single-cell time series classified by experimentalists to construct a clustering strategy to classify cells from future experiments automatically. Singlecell time series for the Cdc10-mCyOFP1 signal were normalized per biological replicate before being pulled together into matrices where each row represented a cell and each column corresponded to a time point. The order of the rows was then randomized and the data was divided into 50 % training data, 30 % validation data, and 20 % test data (Chicco, 2017). The silhouette and the Calinski-Harabasz methods defined the minimum number of clusters in the data, which were confirmed by assessing higher cluster numbers using the function *kmeans()*. Unsupervised clustering algorithms were used to classify cells in the training data set into three clusters according to their Cdc10-mCyOFP1 signal. The algorithms differed in the clustering method (*k*-means, *k*-medoids, hierarchical clustering), the number of time points, the interval of time points, and the cluster centroid distance metric (sqeuclidean, correlation, cityblock, cosine). Clustering algorithms’ performance was evaluated using the validation data, which was manually labeled by three cell biologists tasked with classifying 300 cells from movies as “direct G1 arrest after starvation onset”, “completion of one final cell division after starvation onset” and “persistent arrest as a budded cell after starvation onset”. Clustering algorithms were then asked to classify the human-labeled validation data using the function *pdist2()* and the coherence of the cluster assignments was scored by a multiclass Matthews correlation coefficient (MCC) (Gorodkin, 2004). The best performing classifier algorithms were optimized by adjusting clustering parameters until their clustering closely recapitulated the human cluster assignments as judged by MCC. The centroids of the best clustering solutions were used to write the function *fcentroids()*, which uses the function *pdist2()* and the saved cluster centroids to classify new time series from new experiments into Cdc10-clusters automatically. In strains carrying multiple fluorescent markers, clustering according to Cdc10 was used to arrange all fluorescent channels.

### Clustering analysis of synchronized cultures

For M-phase synchronization, exponentially growing cells, OD_600_ = 0.3, were spun down and resuspended in 20 μg/mL nocodazole-containing SCD and cultured for 2 h (~90 % of dumbbell-shaped cells) before loading in the microfluidics device and being released by exposure to nocodazole-free SCD. For G1 synchronization, an early stationary culture OD_600_ = 2 was centrifuged at 500 rpm for 2 min on a discontinuous PEG 5000: SC gradient (1:3, 1:27, 1:81), and cells in the top 200 ul, which were mostly unbudded, were washed and resuspended in SCD before loading in the flow cell. As soon as 15% of the synchronized population started to bud, cells were transferred to starvation medium.

### Linear discriminant classifier (LDCs) analysis

Affiliation to “Low-Cdk1” or “High-Cdk1” clusters was used as a categorical variable to establish linear discriminant classifiers that measured how well stress and cell cycle markers discriminate between the two clusters at each time point. LDCs hyperparameters were optimized using the function *fitcdiscr()*, holding out 30% of the cross-validation data. Discrimination between clusters at a given time point was measured by the Mahalanobis distance calculated with the function *mahal()*.

### Support Vector Machine (SVMs)-based calculation of single-cell probabilities

Single-cell probabilities of high-Cdk1 quiescence were calculated using a support vector machine (SVM) given by the function *fitcsvm()* with radial basis kernel. SVM’s were trained using the last time point of the starvation period, assuming that stress and cell cycle markers relevant for cluster determination displayed their maximum divergence at this stage. *A priori* probabilities equal to the average ratio of high-Cdk1 cells to the total number of cells were assumed. The trained SVMs were used to classify every cell at every time point according to each stress or cell cycle marker’s values. The function *fitPosterior()* was used to obtain the posterior probability that a cell was correctly assigned to its cluster at a given time point.

### *In-silico* cell sorting according to cellular fluorescence

In copper induction experiments, mTFP1 (+) and control mTFP1 (-) strains were mixed in the same microfluidic space and sorted *in-silico* using the following algorithm (Fig. S4 A): (1) singlecell time series for mTFP1 were transformed into log10 values. (2) the mean of the log10 time series was calculated. (3) The mean log10 values derived from each single-cell time series were clustered using *kmeans()*, distance metric cityblock, which separated mTFP1(-) from MTFP1(+) cells before further analysis.

### Semi-automatic quantifications of single cells and intracellular features

The quantification of the time of SPB pole body separation, time point of cell division, time of maximum nuclear accumulation, and foci detection was made on a previously published pipeline for single-cell analysis with a semi-automated user interface for scoring of single-cell data (Arguello-Miranda et al., 2018), available at https://github.com/alejandrolvido/Quiescence-Entry.

## Online supplemental material

Fig. S1 shows morphological quantifications of quiescent cells under starvation, the algorithm for Cdc10 signal quantification, the workflow of the TPML algorithm, cluster number analysis, and the mitotic induction of the APC/C-Cdh1 activity sensor, *MET3p-mNG-Ase1(R632-I885)*. Fig. S2 shows the assessment of signal-to-noise ratios and crosstalk between fluorescent channels in the six-color imaging setup, the osmoregulatory response in the 6C2 strain, and representative micrographs of cells expressing mNG-Atg8, Q/N-mRuby3, or Rad52-mNG. Fig. S3 shows the clustering analysis of cells treated with nocodazole and nutrient depletion before starvation, as well as the Linear Discriminant analysis for low- and high-Cdk1 Q-cells. Fig. S4 shows the algorithm to sort cells according to the presence/absence of a fluorescent signal, the extended analysis of the *CUP1p*-expression of Xbp1, xbp1-EE, Stb3, and Whi5 during quiescence entry, and the TPML analysis of *xbp1Δ* cells. Fig. S5 shows the analysis of nuclear translocation of stress transcription factors during return to rich medium, cyclic episodes of nutrient depletion, or two consecutive quiescence entries. Video 1 shows the assay to study the proliferation-quiescence transition, including a stress-resistance test by exposure to hyperosmotic medium. Video 2 shows the TPML algorithm being optimized to classify cells in three main clusters. Video 3 shows six-color imaging of the proliferation-quiescence transition in 6C1 cells with single-cell tracking of a representative high-Cdk1 quiescent cell. Video 4 shows six-color imaging of quiescence entry in 6C1 cells *CUP1*p-expressing Xbp1-mTFP1. Video 5 shows six-color imaging of quiescence entry in 6C1 cells *CUP1*p-expressing Whi5-mTFP1. Video 6 shows six-color imaging of quiescence entry in *xbp1Δ* cells bearing cell cycle and stress markers. Video 7 shows six-color imaging of 6C1 cells during cyclic exposure to stress. Video 8 shows six-color imaging of 6C1 cells during two consecutive quiescence entries. Table S1 lists primers used. Table S2 lists plasmids used. Table S3 lists strains used. Table S4 lists the optics set up for six-color imaging.

## Acknowledgments

We thank Gaudenz Danuser, Mike Henne, Sandra Schmid, Jan Skotheim, Jennifer Ewald, and Peter Michaely. This work was supported by the Cancer Prevention and Research Institute of Texas (RP150596 & RR150058), the Welch Foundation (I-1919-20170325), and the National Institute of General Medical Sciences of the National Institutes of Health (K99GM135487). OA conceived the idea, analyzed data, and wrote computer code. JN designed statistical analyses. JN and OA wrote the paper. AM, TK, MR, and OA performed experiments. We thank Andreas Doncic posthumously.

