## Supplemental Material for "Cell cycle-independent integration of stress signals promotes Non-G1/G0 quiescence entry"

### Supplemental Information

**Fig. S1 Analysis of single cell morphological features during quiescence entry, Cdc10 signal quantification algorithm workflow, TPML algorithm workflow, cluster number analysis, and mitotic induction of the APC/C-Cdh1 activity sensor, *MET3p-mNG-Ase1(R632-I885)*. (A)** Comparison of morphologic parameters between budded and unbudded quiescent cells. **(B)** Algorithm to detect cell cycle progression based on the standard deviation of Cdc10-mCyOFP1 fluorescence at the cell periphery or "Cdc10 Signal a.u.". **(C)** Generation of an ML-algorithm to sort cells into clusters depending on the pattern of cell cycle arrest indicated by Cdc10. (Left) Representative single-cell time series for Cdc10-mCyOFP1 signal (*OAM421)*. ***(1)*** time series are normalized per biological replicate and their cell index randomized. ***(2)*** time series data are split into 50% training time series, 30% validation time series, and 20% test time series. ***(3)*** Training time series are used to establish different clustering models depending on parameters such as clustering method (k-means, k-medoids), time window (number of time points in time series), number of clusters, and distance metric. ***(4)*** Validation time series are labeled by experienced cell biologists who assign different cluster affiliations to each time series manually. ***(5)*** the preliminary clustering models are optimized by comparing the unsupervised clustering solutions to the human clustering solution using a multiclass Matthews's coefficient. ***(6a)*** the best performing clustering model is tested using the ***(6b)*** test time series. The best performing algorithm is used to classify cells in future experiments. **(D)** Evaluation of the number of clusters using the silhouette method or by inspecting increasing number of clusters (*k* values). *k*=3 is the minimum number of clusters that recapitulate the major cell cycle arrest patterns in the population. **(E)** Induction of the APC/C-Cdh1 activity sensor, mNG-Ase1(R632-I885), upon depletion of methionine in rich medium (green arrow); *OAM487* cells were aligned *in silico* according to their Cdc10-signal and the average times series was plotted, (right) Cdc10 (left) APC/C-Cdh1 sensor. Solid lines with shaded area = average ± 95% confidence intervals.

**Fig. S2 Analysis of signal-to-noise ratios and bleed-through in the six-color imaging system, analysis of osmoregulatory responses in 6C2 cells, and detection of mNG-Atg8, Q/N-mRuby3, or Rad52 foci. (A-H)** Assessment of signal-to-noise ratios and bleed-through in a six-color fluorescent microscopy system by imaging quiescence entry in isogenic single fluorophore-tagged stains for **(A)** Cdc10-mCyOFP1 (*OAM421)* and the stress markers **(B)** Sfp1-mScarlet-I (*OAM407)*, **(C)** Gln3-mKO*κ (OAM507)*, **(D)** Rtg1-mNeptune2.5 (*OAM510*), **(E)** Stb3-mTFP1 (*OAM395*), **(F)** Xbp1-mNeonGreen (*OAM397*) and **(G)** isogenic blank cells (*OAM128*) or **(H)** an isogenic six-color strain (*OAM425*); at least 75 cells per strain were analyzed from 3 different biological replicates; each channel is normalized to a mean of 100 a.u. Plot with colored background = channel containing a fluorescent signal. Plot with white plot background = channel without a fluorescent signal. Red arrow = onset of starvation. **(I)** Upregulation of osmotic/mechano-sensitive stress responses by 200 mM $CaCl_2$ in unsynchronized 6C2 cells (*OAM667*) bearing the stress markers Crz1-mTFP1, Pkc1-mRuby3, Msn2-mNeonGreen, Sfp1-mKOκ, Hog1-mNeptune2.5 and Cdc10-mCyOFP1. Heatmaps display unsynchronized single-cell time series. The plots display average time series (n=4, 364 cells). Red area, period in 200mM $CaCl_2$-containing rich medium. Green triangle, onset of $CaCl_2$ exposure. **(J)** Representative micrographs showing the accumulation of the autophagy reporter mNeonGreen-Atg8 upon exposure to low nitrogen medium (without ammonium sulfate) during proliferation (*OAM654*) **(K)** Representative time-lapse micrographs of a mother cell retaining a fluorescent QN-mRuby3 aggregate during proliferation in rich medium after a 2 h in 25 μM $CuSO_4$-containing SCD (*OAM671)*. **(L)** Representative time-lapse micrographs of cells (*OAM670*) with recurring Rad52-mNeonGreen foci during starvation. **(M)** Average single cell QN-mRuby3 foci intensity during quiescence entry per Cdc10 Cluster. **(N)** Average number of Rad52-mNeonGreen foci during quiescence entry per Cdc10 Cluster. Solid lines with shaded area=average ± 95% confidence intervals.

**Fig. S3 Analysis of quiescence entry after cell cycle-synchronization or nutrient depletion and linear discriminant analysis of cluster separation by stress factors. (A)** a 2-step *k*-means-based algorithm to sort time series before TPML. Heatmaps = single-cell time series derived from 6C1 cells that were M-phase (nocodazole) synchronized before starvation. Clustering identifies cells that exited from G1 shortly after starvation onset. **(B)** Average time series for cell cycle (Cdc10 signal) and stress markers (Sfp1, Rtg1, Gln3, Stb3, Xbp1) in M-phase arrest-released 6C1 cells that exited G1 shortly before starvation onset (n=3, 96 cells *OAM425*). **(C)** Average time series for cell cycle (Cdc10 signal) and stress markers (Sfp1, Rtg1, Gln3, Stb3, Xbp1) in G1-released 6C1 cells that exited G1 shortly before starvation onset (n=3, 86 cells *OAM425*). **(D)** Average time series of cell cycle (Cdc10 signal) and stress markers (Sfp1, Rtg1, Gln3, Stb3, Xbp1) in 6C1 cells that were exposed to glucose depletion for 2 h before quiescence entry (n=3, 504 cells *OAM425*). **(E)** Average time series of cell cycle (Cdc10 signal) and stress markers (Sfp1, Rtg1, Gln3, Stb3, Xbp1) in 6C1 cells that were exposed to nitrogen depletion for 2 hours before quiescence entry (n=3, 222 cells *OAM425*). **(F)** Average cluster separation at each time point between low- and high-Cdk1 Q-cells, as measured by the Mahalanobis distance calculated using linear discriminant classifiers (LDC) based on the time series for each stress marker derived from *OAM425* (Fig. 3) and *OAM667* (Fig. 4) cells. Solid lines with shaded area = average ± 95% confidence intervals from biological replicates. Green arrow = onset of nutrient depletion. Red Arrow = onset of starvation.

**Fig. S4 Sorting of time series according to the presence/absence of a fluorescent signal, analysis of quiescence entry in cells *CUP1*p-expressing transcriptional repressors, and in *xbp1*Δ cells.** **(A)** Sorting of mixed time series from cells expressing mTFP1-tagged transcriptions factors and control cells without the mTFP1 construct that were loaded in the same microfluidics device (See figure 7). **(B-E)** Analysis of quiescence entry in strains bearing Gln3-mKOκ, Rtg1-mNeptune2.5, Sfp1-mScarlet-I, Xbp1-mNG and Cdc10-mCyOFP1 after *CUP1*p-expression of mTFP1 C-terminally tagged wild-type Xbp1 (n=6 *OAM458*), the DNA binding domain mutant xbp1-EE (n=7 *OAM466*), the Xbp1-related transcriptional repressor Stb3 (n=4 *OAM497*) or the cell cycle repressor Whi5 (n=7 *OAM455*) at starvation onset. **(B)** Average time series per cluster for cell cycle (Cdc10 signal) and stress markers (Sfp1, Rtg1, Gln3, Xbp1) in cells expressing *CUP1p-XBP1-mTFP1* during quiescence entry. Cells whose Xbp1-mTFP1 levels exceeded two standard deviations from the mean Xbp1-mTFP1 levels were excluded. **(C)** Average time series per cluster for cell cycle (Cdc10 signal) and stress markers (Sfp1, Rtg1, Gln3, Xbp1) in cells expressing *CUP1p-xbp1-EE-mTFP1* during quiescence entry. **(D)** Average time series per cluster for cell cycle (Cdc10 signal) and stress markers (Sfp1, Rtg1, Gln3, Xbp1) in cells expressing *CUP1p-STB3-mTFP1* during quiescence entry. **(E)** Average time series per cluster for cell cycle (Cdc10 signal) and stress markers (Sfp1, Rtg1, Gln3, Xbp1) in cells expressing *CUP1p-WHI5-mTFP1* during quiescence entry. **(F) top,** Average time series per cluster for cell cycle (Cdc10 signal) and stress markers (Sfp1-mScarlet-I, Gln3-mKOκ, Rtg1-mNeptune2.5, Msn2-mNG and Stb3-mTFP1) in six-color *xbp1*Δ cells (n=4, 500 cells *OAM500).* Unlike *XBP1* strains, quiescence entry in *xbp1*Δ cells could not be described as three main clusters; instead, TPML sorted *xbp1*Δ cells into four clusters, indicated by the 4-color bar on the left on the bottom heat maps. Gray area on plot = period in rich medium. Solid lines with shaded area = average ± 95% confidence intervals.

**Fig. S5 Analysis of nuclear translocation of stress transcription factors upon return to rich medium and during cyclic episodes of glucose/nitrogen depletion or two consecutive quiescence entries.** **(A)** Average time series for cell cycle (Cdc10 signal) and stress markers (Sfp1, Rtg1, Gln3, Stb3, Xbp1) during return to proliferation in 6C1 cells (n=6, *OAM425*); green dotted line, transfer from starvation to rich medium. **(B)** Nuclear intensity of Sfp1, Stb3, Xbp1, Rtg1, Gln3, and cell size, expressed as percentage of their values at 20 h of starvation, during cell divisions upon return to rich medium (n=4, > 101 cells *OAM425*). **(C)** Average times series for cell cycle (Cdc10-mCyOFP1) or nuclear stress makers (Sfp1, Gln3, Rtg1, Stb3, Xbp1) during hourly cycles of glucose depletion (n=5, 335 cells *OAM425*). **(D)** Average times series for cell cycle (Cdc10-mCyOFP1) or nuclear stress makers (Sfp1, Gln3, Rtg1, Stb3, Xbp1) during hourly cycles of nitrogen depletion (n=5, 235 cells *OAM425*). **(E)** Average nuclear Stb3-mTFP1 intensity in cells transferred to rich medium after different starvation periods (n=3, >71 cells *OAM395*). **(F)** Average of Cdc10-clusters for cell cycle (Cdc10-mCyOFP1) and stress markers (Sfp1, Stb3, Xbp1, Gln3, and Rtg1) during two consecutive quiescence entries (n=3, 354 cells). Cdc10 clusters were established according to the first quiescence entry; only cells present during the first quiescence entry are included. **(G)** Percentage of high-Cdk1 Q-cells during two consecutive quiescence entries in control (n=3, 300 cells) or *xbp1*Δ cells (n=4, 450 cells). Gray rectangles, period in rich medium. Solid lines with shaded area = average ± 95% confidence intervals. Red star, $p<0.05$, KS-test. Boxplots display data from biological replicates: central mark, median; box bottom and top limit, 25 th and 75 th percentiles; whiskers, most extreme non-outlier values.

#### Supplemental Movie Legends

**Video 1.** A microfluidics assay to study the transition from proliferation-into-quiescence. After 90 min in rich medium, proliferating Wild-Type cells (OAM128) are exposed to starvation medium for 20 h and reached a stress-resistant quiescent state that survives exposure to 4 M NaCl for 4 h before resumption of proliferation by exposure to rich medium.

**Video 2.** Time series profiling by machine learning algorithm (TPML) training on single-cell time series (Cdc10-mCyOFP1 data). In this example, TPML finds the best interval (number of time points) for clustering cells according to cell cycle arrest pattern. Each frame corresponds to a potential clustering solution evaluated against a human-labeled validation data set using the Matthews correlation coefficient (MCC). Clustering solutions with the highest MCC are further optimized and eventually converge in a clustering solution (See Fig. S1B-C).

**Video 3.** Six-color imaging of the proliferation-quiescence transition in 6C1 cells (*OAM425*) carrying the cell cycle marker Cdc10-mCyOFP1 and the stress response markers Sfp1-mScarlet-I, Gln3-mKO*κ,* Rtg1-mNeptune2.5, Stb3-mTFP1 and Xbp1-mNeonGreen. Each peripheral hexagon displays a median-filtered fluorescent channel. Each subplot corresponds to a time series obtained from the representative high-Cdk1 Q-cell highlighted in yellow.

**Video 4**. Six-color imaging of cells *CUP1*p-expressing Xbp1-mTFP1 (*OAM458)* and control cells (without Xbp1-mTFP1, *OAM454*) quiescence entry in. Unlike control cells, *CUP1-XBP1-mTFP1* cells underwent high-Cdk1 quiescence entry with high frequency. Notice the brighter Xbp1-mNeonGreen nuclear signal in control cells. Cells also carry the cell cycle marker Cdc10-mCyOFP1 and the stress response markers Sfp1-mScarlet-I, Gln3-mKO*κ,* Rtg1-mNeptune2.5, and Xbp1-mNeonGreen. Each peripheral hexagon displays a median-filtered fluorescent channel.

**Video 5**.Six-color imaging of cells *CUP1*p-expressing Whi5-mTFP1 (*OAM455)* during quiescence entry*.* Notice that high-Cdk1 Q-cells expressing *CUP1p-WHI5-mTFP1* keep Whi5-mTFP1 in the cytoplasm during starvation (budded cell in the middle). Cells also carry the cell cycle marker Cdc10-mCyOFP1 and the stress response markers Sfp1-mScarlet-I, Gln3-mKO*κ,* Rtg1-mNeptune2.5, and Xbp1-mNeonGreen. Each peripheral hexagon displays a median-filtered fluorescent channel.

**Video 6.** Six-color imaging of quiescence entry in *xbp1Δ* cells (*OAM500*). Cells carry Cdc10-mCyOFP1 as cell cycle marker and Sfp1-mScarlet-I, Gln3-mKO*κ,* Rtg1-mNeptune2.5, Stb3-mTFP1 and Msn2-mNeonGreen, as stress response markers. Notice how *xbp1Δ* cells fail to arrest after starvation. Each peripheral hexagon displays a median-filtered fluorescent channel.

**Video 7.** Six-color imaging of 6C1 cells (*OAM425*) during cyclic exposure to stress episodes triggered by hourly changes between rich (SCD) and glucose-deficient medium (SCG). Cells carry Cdc10-mCyOFP1 as cell cycle marker and Sfp1-mScarlet-I, Gln3-mKO*κ,* Rtg1-mNeptune2.5, Stb3-mTFP1 and Xbp1-mNeonGreen, as stress response markers. Each peripheral hexagon displays a median-filtered fluorescent channel.

**Video 8.** Six-color imaging of 6C1 cells (*OAM425*) during two consecutive quiescence entries separated by a period in rich medium. Cells carry Cdc10-mCyOFP1 as cell cycle marker and the stress response markers Sfp1-mScarlet-I, Gln3-mKO*κ,* Rtg1-mNeptune2.5, Stb3-mTFP1 and Xbp1-mNeonGreen. Each peripheral hexagon displays a median-filtered fluorescent channel.

### Supplemental Figure 1

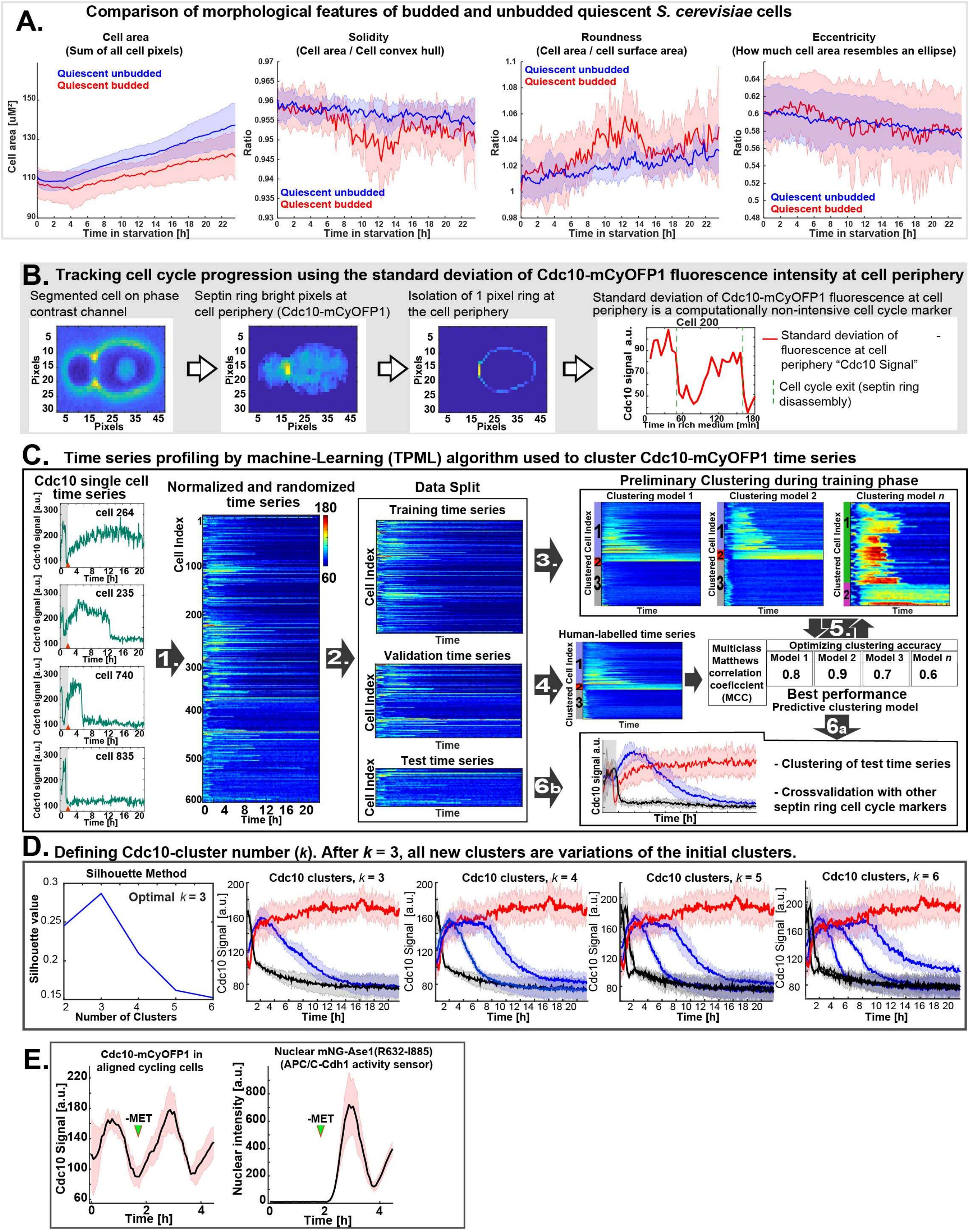

### Supplemental Figure 2

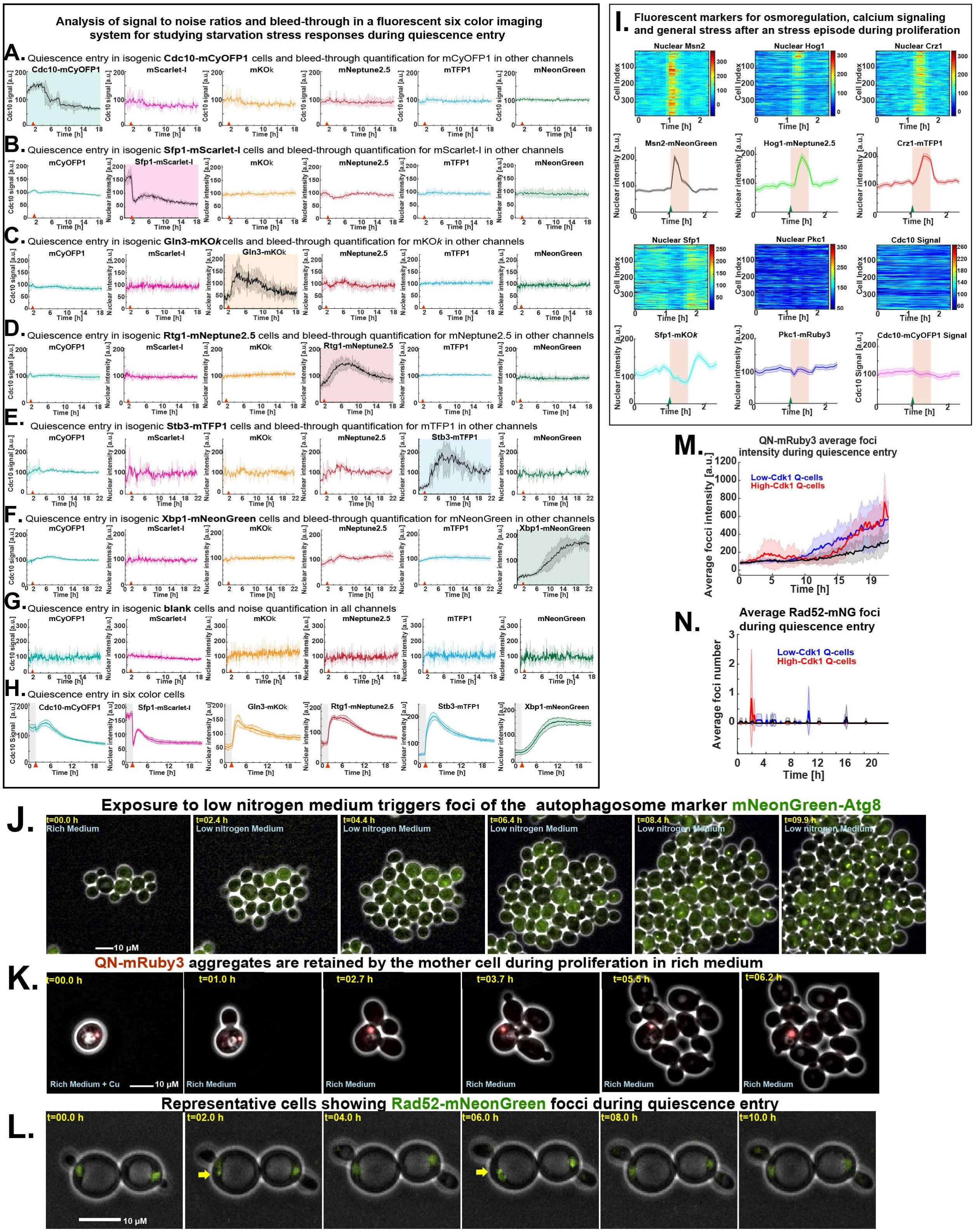

### Supplemental Figure 3

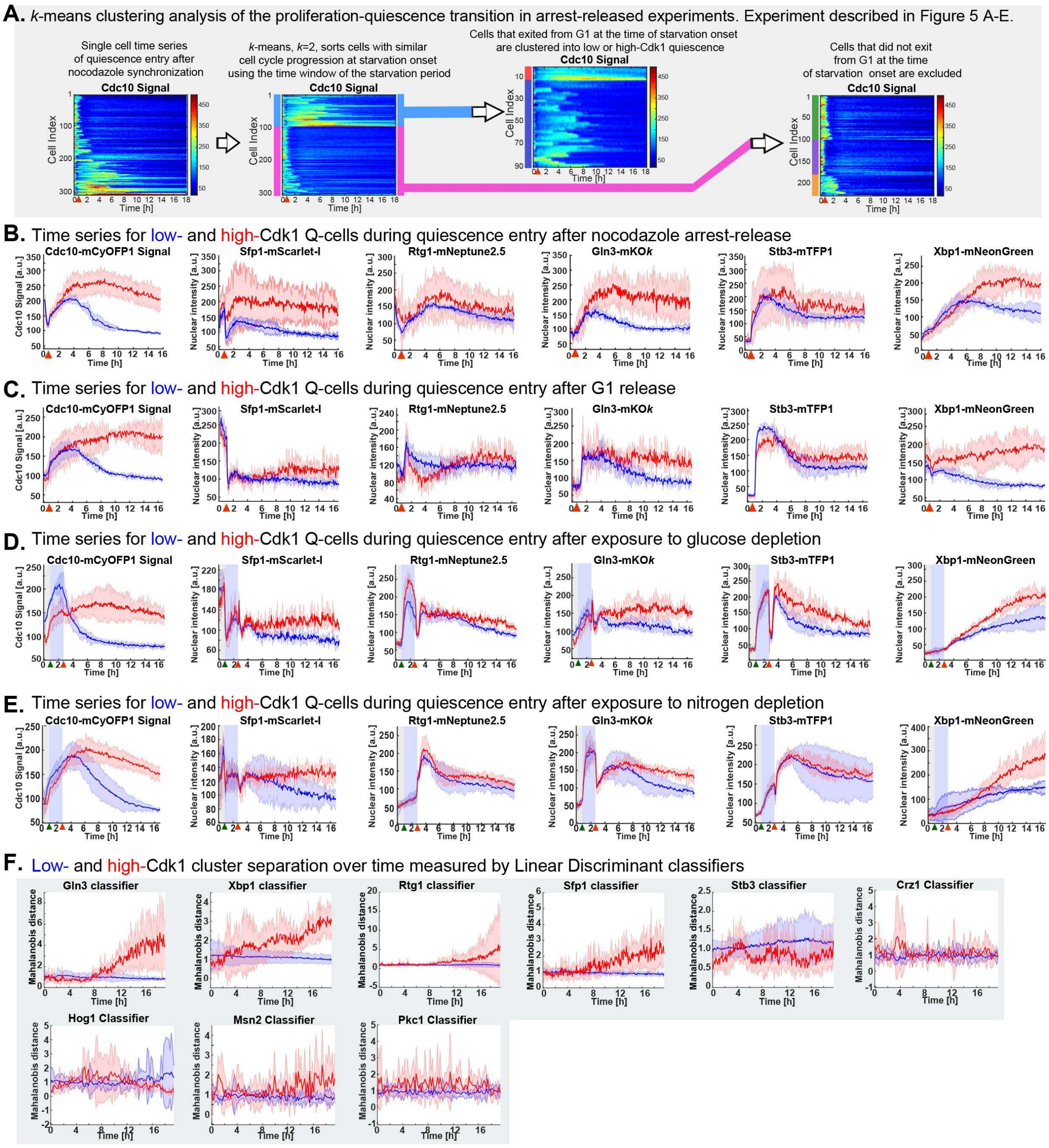

### Supplemental Figure 4

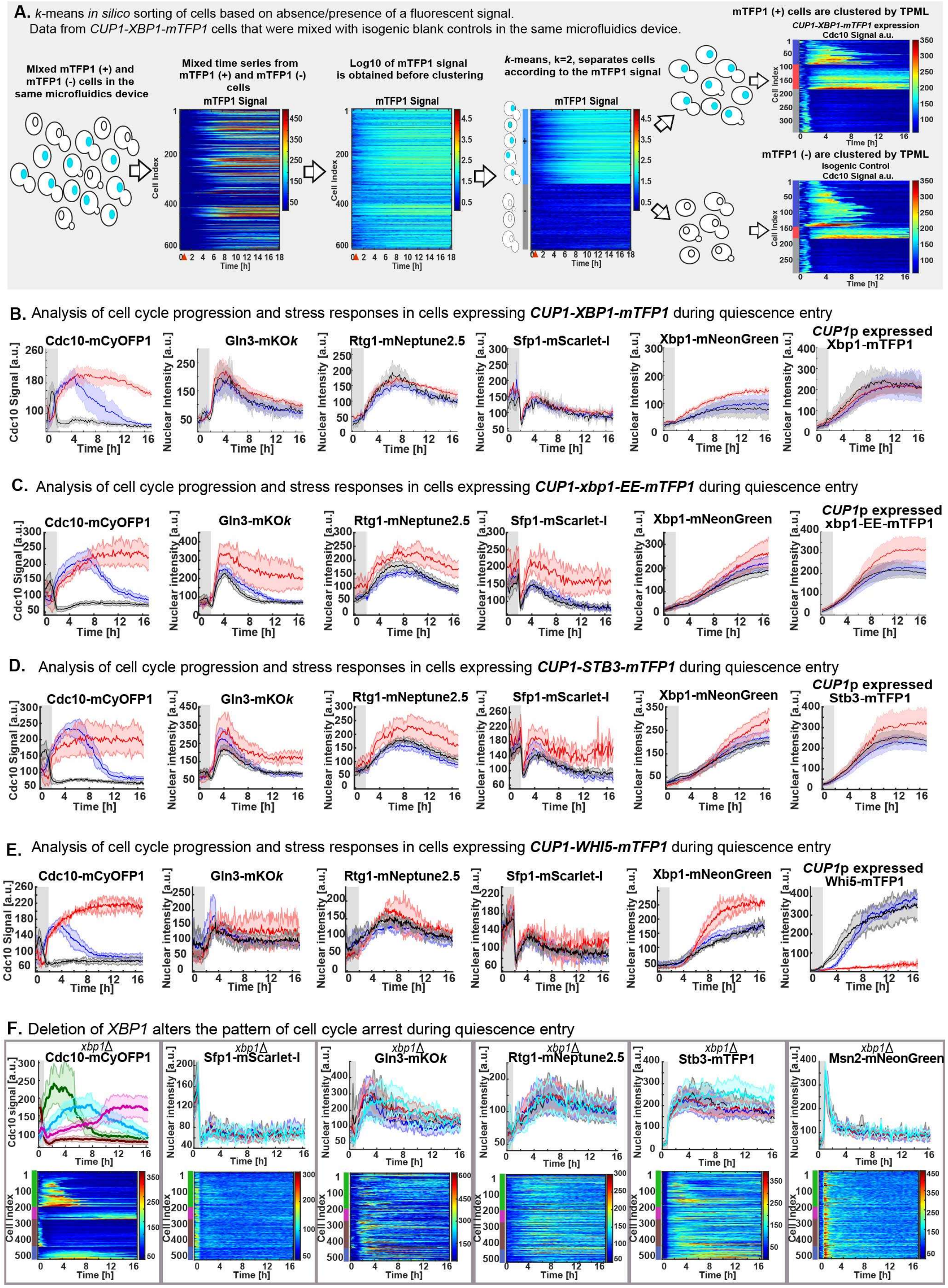

### Supplemental Figure 5

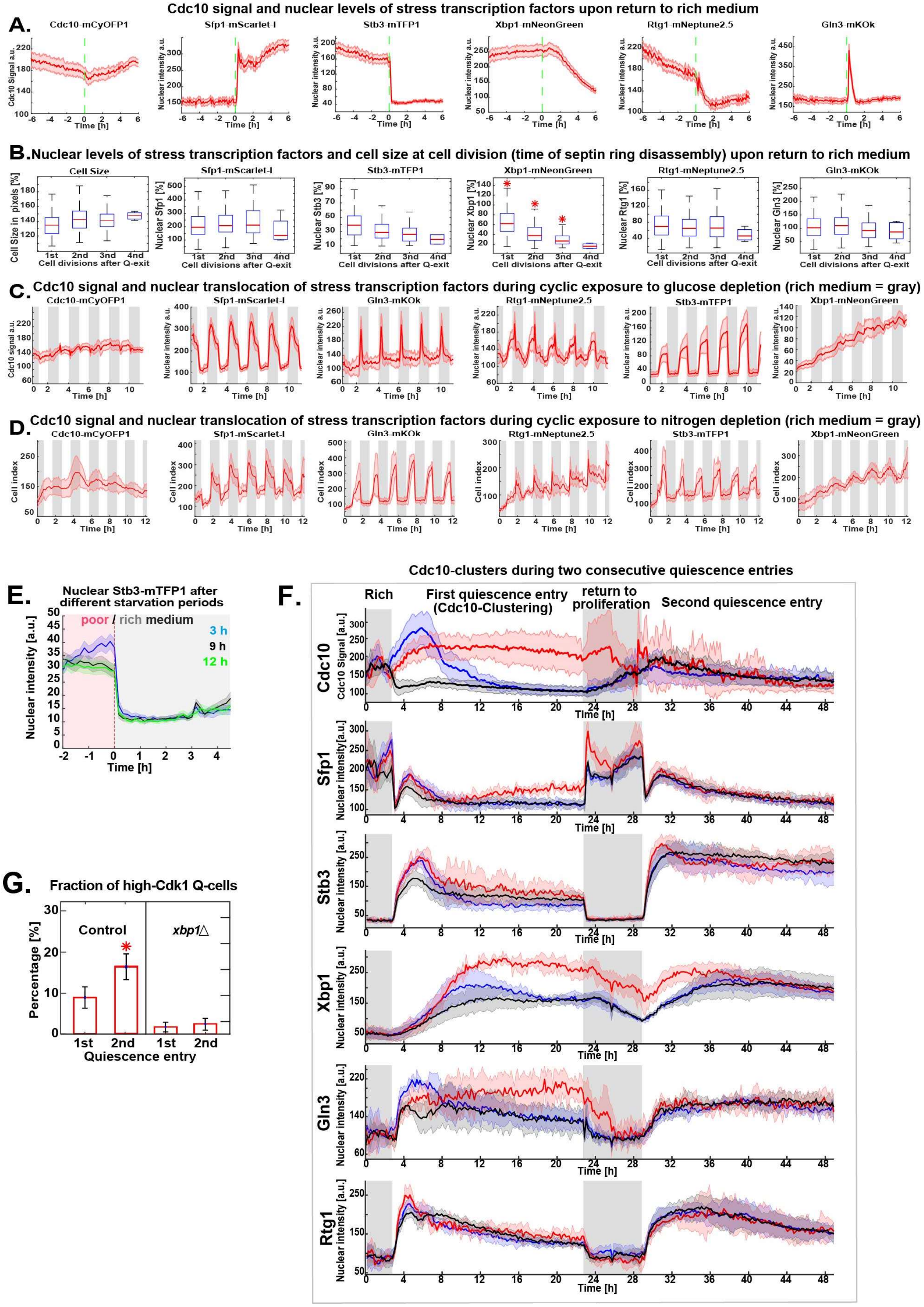

### Supplemental Tables

**Supplementary Table 1. Primers used for strain construction.**

| Oligonucleotide | 5'-Sequence-3' |
|---|---|
| Cdc10-F | TCGTTCCTCAGCTCATATGTCTAGCAACGCCATTCAACGTCGGATCCCCGG GTTAATTAA |
| Cdc10-R | AATAACATAAGATATATAATCACCACCATTCTTATGAGATGAATTCGAGCTCG TTTAAAC |
| Sic1-F | GAGAAGATTCAAGCCAAAGGCATTGTTTCAATCTAGGGATCAAGAGCATCG GATCCCCGGGTTAATTAA |
| Sic1-R | GTAAGTAAATAAAATATAATCGTTCCAGAAACTTTTTTTTTCATTTCTGAATT CGAGCTCGTTTAAAC |
| Ase1-AscI-F | TTTAGAAGTGGCGCGCCTCAGATCTAAAAAGGGAAAATG |
| Ase1-NotI-R | CTGATTGCGGCCGCTCAAATATCTGTAAAGGAGAATCC |
| Met3-SacI-F | TATCAGAGCTCTTTAGTACTAACAGAGAC |
| Met3-PacI-R-new | CTGATTTTAATTAAGTTAATTATACTTTATTCTTG |
| mNeonGreen-BamHI | ATAACAACAGGATCCCTTGTACAGCTCGTCCATGCCC |
| mNeonGreen-PacI | ATAACAACATTAATTAACATGGTGAGCAAGGGC |
| Cdc7-R | AAAGGAAATTATTTCAGACTAGCAGTAATTTATCACTTTGGAATTCGAGCTCG TTTAAAC |
| Cdc7-F | GGCAGATTTGCTCGATAAGGATGTTCTCCTAATATCTGAACGGATCCCCGG GTTAATTAA |
| Spc42-F | TATGTCAGAAACATTCGCAACTCCCACTCCCAATAATCGACGGATCCCCGG GTTAATTAA |
| Spc42-R | TTTAAGAATGCGCCATACTCCTTAACTGCTTTTTAAATCAGAATTCGAGCTCG TTTAAAC |
| Cdc14-F | TTCAAAAAAATAATATCTTCATTCAATCATGATTCTTTTTGAATTCGAGCTCGT TTAAAC |
| Cdc14-R | CGGTGCTGGTCACTCCAACACTTTGCAAGTCTCTACCGTTCGGATCCCCGG GTTAATTAA |
| Gln3-F | AGCAATTGCTGACGAATTGGATTGGTTAAAATTTGGTATACGGATCCCCGGG TTAATTAA |
| Gln3-R | TTATTAACATAATAAGAATAATGATAATGATAATACGCGGGAATTCGAGCTCG TTTAAAC |
| Rtg1-F | CTTCGAGTACGGAGGGTATGGTGAGTACGGTAATGGTAGCCGGATCCCCG GGTTAATTAA |
| Rtg1-R | GGTTATCACAACATAGCAATAGTGAGAGTCAGAAGTACTTGAATTCGAGCTC GTTTAAAC |
| Stb3-F | AGACAATAGTGCTGCGTTTTTATTAATGAGCTTAAAATCTCGGATCCCCGGG TTAATTAA |
| Stb3-R | TACTGTTTTTTTGTTATTTCATGGAACTGGTAAAGAATTCGAATTCGAGCTCG TTTAAAC |
| Xbp1-F | GGGCGAACAATTCACTGAACAATTTCAAATTTAAAACAAACTCAAAACAACG GATCCCCGGGTTAATTAA |

| | |
|---|---|
| Xbp1-R | AAAAATTAAAGAACCATTATAAAAGAATATTACCAAAAATGTAACAGATGACA TTTGAATTCGAGCTCGTTTAAAC |
| Sfp1-F | GTGGTAAGAGATATAAGAACTTGAACGGTTTAAAATATCACAGGGGCCACTC CACTCACCGGATCCCCGGGTTAATTAA |
| Sfp1-R | CGGTAGAATTAGTATCTAAATAGCATACTAAATCAATTCAGTAAAGAAACAAT TATATCGAATTCGAGCTCGTTTAAAC |
| Crz1-F | CATCACTCCCTTGTACGAAGAAGCCAGACAGGAGAAATCGGGACAAGAGAG TCGGATCCCCGGGTTAATTAA |
| Crz1-R | CTACACTGTCATTATGAAATTACATATTTATTATATAGAAAAAAAAAATTCCTA TTCAAAGCTTAAAAAAACAAAAATAAGAATTCGAGCTCGTTTAAAC |
| Hog1-F | CGGTAACCAGGCCATACAGTACGCTAATGAGTTCCAACAGCGGATCCCCGG GTTAATTAA |
| Hog1-R | GAAGTAAGAATGAGTGGTTAGGGACATTAAAAAAACACGTGAATTCGAGCTC GTTTAAAC |
| Pkc1-F | GACAACGAGCCAGCAAGAAGAGTTTAGAGGATTTTCCTTTATGCCAGATGAT TTGGATTTA CGGATCCCCGGGTTAATTAA |
| Pkc1-R | GCGGAACAGTTCAACTTTTCCGCTTAGATGTTTTATATAAAATTAAATAAATC ATGGCATGACCTTTTCTGAATTCGAGCTCGTTTAAAC |
| Msn2-F | GTCGCAACACATCAAGACTCATAAAAAACATGGAGACATTCGGATCCCCGG GTTAATTAA |
| Msn2-R | TTATGAAGAAAGATCTATCGAATTCCCCCCCTGGGGTCTAGAATTCGAGCTC GTTTAAAC |
| ATG8-ORF-R | CACAGGTATCCTATTCTTGAAC |
| ATG8-ORF-F | GTTCAAGAATAGGATACCTGTG |
| QNgln3-HindIII-F | TAATAAGCTTATGTCTCAATACAACCACGGTTCC |
| QNgln3-PacI-R | CTGATTTTAATTAACTGGATATTACTATTGTTGCT |
| Cup1-SalI-F | GCTGTCGACCTATACGTGCATATGTTCATG |
| Cup1-HindIII-R | CGTAAGCTTTTGATTGATTGTACAGTTTG |
| Cup1-NheI-R | TGTTGTTATGCTAGCTTGATTGATTGTACAGTTTG |
| Tom70-F | GGCTAAAAAGATTCAAGAAACTTTAGCTAAATTACGCGAACAGGGTTTAATG CGGATCCCCGGGTTAATTAA |
| Tom70-R | CTACTTTGTTACTTAGTTTTTGTCTTCTCCTAAAAGTTTTTAAGTTTATGTTTA CTGTGAATTCGAGCTCGTTTAAAC |
| Rad52-F | GCCAGCCGTTCCGCAACAAAGATCGACACGAAGAGAAGTTGGAAGACCAAA GATCAATCCCCTGCATGCACGCAAGCCTACTCGGATCCCCGGGTTAATTAA |
| Rad52-R | CAGAATTGAAAGAGTAACTAGAGGATTTTGGAGTAATAAATAATGATGCAAAT TTTTTATTTGTTTCGGCCAGGAAGCGTTGAATTCGAGCTCGTTTAAAC |
| Xbp1-HindIII-F | TAATAAGCTTTATATGAAATATCCCGCTTTTAGC |
| Xbp1-PacI-R | CTGATTTTAATTAATTGTTTTGAGTTTGTTTTAAATTTG |
| Whi5-HindIII-F | TAATAAGCTTTATATGAGTTTGAGAACGCCG |
| Whi5-PacI-R | CTGATTTTAATTAAAGACGTCTCCACTTCGGTATCCG |
| Stb3-HindIII-F | ATACACAAAGCTTATGTCAGAAAACCAAAAGGAGG |
| Stb3-PacI-R | ATACACATTAATTAAAGATTTTAAGCTCATTAATAAAAACGC |
| Xbp1-ClaI-DBD-F | CAATTACATCGATTTTCATTGG |

| | |
|---|---|
| Xbp1-DBD-E-R | CCTTGAGATTTCCATCGGCAACCAAGTACCCTCAATTTTGATGTAACCACCT TCTATTCTTTTC |
| Xbp1-del-F | CATATATGAATGCCGGGCCG |
| Xbp1-del-R | CTTCGGCACGTCATACACC |

**Supplementary Table 2. Plasmids used for strain construction.**

| Plasmid ID | Fluorophore | Marker | Construction |
|---|---|---|---|
| pOAM0 | mTFP1 | URA3 | The mCherry fluorophore in pLondon266-mCherry was cut out with *Pac*I and *Asc*I and replaced with an mTFP1 fluorophore |
| pOAM1 | mTFP1 | URA3 | Triple ligation of (1) *XBP1* flanked by *Hind*III and *Pac*I restriction sites, (2) *CUP1* promoter (434-16 bp upstream of start codon) flanked by *Sal*I and *Hind*III restriction sites, and (3) pLondon266-mTFP1 vector cut with *Sal*I and *Pac*I |
| pOAM2 | mTFP1 | URA3 | Triple ligation of (1) *WHI5* flanked by *Hind*III and *Pac*I restriction sites, (2) *CUP1* promoter flanked by *Sal*I and *Hind*III restriction sites, and (3) pLondon266-mTFP1 vector cut with *Sal*I and *Pac*I |
| pOAM3 | mTFP1 | URA3 | Triple ligation of (1) *STB3* flanked by *Hind*III and *Pac*I restriction sites, (2) *CUP1* promoter flanked by *Sal*I and *Hind*III restriction sites, and (3) pLondon266-mTFP1 vector cut with *Sal*I and *Pac*I |
| pOAM6 | mTFP1 | URA3 | Site-directed mutagenesis changed both R349 and Q356 to E in the *XBP1* DNA binding domain. pOAM1 was cut with *Cla*I and *Bsa*BI and was replaced by the EE mutant sequence |
| pOAM7 | mNeonGreen (Yeast Optimized) | LEU2 | Quadruple ligation of (1) *ASE1(aa632-885)* fragment flanked by *AscI* and *Not*I restriction sites (2) *MET3* promoter flanked by *Sac*I and *Pac*I restriction sites, (3) mNeonGreen flanked by *Pac*I and *Asc*I restriction sites, and (4) pRS305 vector cut with *Sac*I and *Not*I. |
| pOAM7 | ATG8p-mNeonGreen (Yeast Optimized)-ATG8 | LEU2 | A pRS304 plasmid carrying the autophagy sensor *ATG8p-mNeonGreen-ATG8-ATG8ter* was created by replacing the *Pac*I-*Bam*HI flanked 2xyeGFP sequence in a pRS304-ATG8p-2xyeGFP-ATG8 (Henne Lab) with a *Pac*I-*Bam*HI-flanked mNeonGreen fluorophore |
| pOAM8 | mRuby3 | URA3 | Triple ligation of (1) a SalI-*Hind*III flanked *CUP1* promoter sequence (2) a *Hind*III-*Pac*I flanked glutamine and asparagine-rich sequence of *GLN3* (aa166-242) (3) *Sal*I-*Pac*I pLondon266 backbone containing the mRuby3 fluorophore. |
| pOAM9 | mTFP1 | HIS3 | pLongtine-Citrine-His was cut with *Pac*I and *Asc*I and the Citrine fluorophore was replaced by mTFP1 flanked by *Pac*I and *Asc*I restriction sites |

| | | | |
|---|---|---|---|
| pOAM10 | mKOκ | HphMX6 | pCA14 was cut with *Pac*I and *Asc*I and the GFP fluorophore was replaced by mKOκ flanked by *Pac*I and *Asc*I restriction sites |
| pOAM11 | mScarlet-I | KanMX4 | pLongtine4 was cut with *Pac*I and *Asc*I and the GFP fluorophore was replaced by mScarlet-I flanked by *Pac*I and *Asc*I restriction sites |
| pOAM12 | mScarlet-I | HphMX6 | pCA14 was cut with *Pac*I and *Asc*I and the GFP fluorophore was replaced by mScarlet-I flanked by *Pac*I and *Asc*I restriction sites |
| pOAM13 | mNeonGreen (Yeast Optimized) | HphMX6 | pCA14 was cut with *Pac*I and *Asc*I and the GFP fluorophore was replaced by mNeonGreen flanked by *Pac*I and *Asc*I restriction sites |
| pOAM14 | mCyOFP1 | NatMX6 | pCA13 was cut with *Pac*I and *Asc*I and the GFP fluorophore was replaced by mCyOFP1 flanked by *Pac*I and *Asc*I restriction sites |

**Supplementary Table 3. Strain list.** *Saccharomyces Cerevisiae* diploid strains are isogenic to W303 (*leu2-3,112 his3-11,15 ura3-1 trp1-1 can1-100 ade2*) and carry heterozygous fluorescent markers unless otherwise stated.

| Strain | Genotype |
|---|---|
| *OAM128* | *MATa/MATα, HIS3/his3, TRP1/trp1, LEU2/leu2, URA3/ura3* |
| *OAM394* | *MATa/MATα, HIS3/his3, TRP1/trp1, LEU2/leu2, URA3/ura3, TPS1/TPS1-mNeonGreen-hphMX* |
| *OAM396* | *MAT a/MATα, HIS3/his3, TRP1/trp1, LEU2/leu2, URA3/ura3, GSY2/GSY2-mNeonGreen-hphMX* |
| *OAM421* | *MATa/MATα, HIS3/his3, TRP1/trp1, LEU2/leu2, URA3/ura3, CDC10/CDC10-mCyOFP1-natMX* |
| *OAM672* | *MATa/MATα, HIS3/his3, TRP1/trp1, LEU2/leu2, URA3/ura3, SIC1/SIC1-mNeonGreen-hphMX, CDC10/CDC10-mCyOFP1-natMX.* |
| *OAM487* | *MATa/MATα, HIS3/his3, TRP1/trp1, leu2/ leu2::MET3p-mNeonGreen-ASE1(aa632-885)-LEU2, URA3/ura3, CDC10/CDC10-mCyOFP1-natMX* |
| *OAM423* | *MATa/MATα, HIS3/his3, TRP1/trp1, LEU2/leu2, URA3/ura3, SPC42/SPC42-mTFP-HIS3, CDC10/CDC10-mCyOFP1-natMX* |
| *OAM422* | *MATa/MATα, HIS3/his3, TRP1/trp1, LEU2/leu2, URA3/ura3, CDC7-mScarlet-I-hphMX/CDC7-mScarlet-I-hphMX, CDC10/CDC10-mCyOFP1-natMX* |
| *OAM407* | *MATa/MATα, HIS3/his3, TRP1/trp1, LEU2/leu2, URA3/ura3, SFP1/SFP1-mScarlet-I-hphMX* |
| *OAM507* | *MATa/MATα, HIS3/his3, TRP1/trp1, LEU2/leu2, URA3/ura3, GLN3/GLN3-mKOκ-hphMX* |
| *OAM397* | *MATa/MATα, HIS3/his3, TRP1/trp1, LEU2/leu2, URA3/ura3, XBP1/XBP1-mNeonGreen-hphMX* |
| *OAM395* | *MATa/MATα, his3/his3, TRP1/trp1, LEU2/leu2, URA3/ura3, STB3/STB3-mTFP1-HIS3* |
| *OAM510* | *MATa/MATα, HIS3/his3, TRP1/trp1, LEU2/leu2, URA3/ura3, RTG1-mNeptune2.5-kanMX/RTG1-mNeptune2.5-kanMX* |

| | |
|---|---|
| *OAM425* | *MATa/MATα, his3/his3, TRP1/trp1, LEU2/leu2, URA3/ura3, STB3/STB3-mTFP1-HIS3, RTG1-mNeptune2.5-kanMX4 /RTG1-mNeptune2.5-kanMX4, SFP1-mScarlet-I-hphMX /SFP1-mScarlet-I-hphMX, GLN3/GLN3-mKOκ-hphMX, XBP1/XBP1-mNeonGreen-hphMX, CDC10/CDC10-mCyOFP1-natMX* |
| *OAM667* | *MATa/MATα, HIS3/his3, TRP1/trp1, LEU2/leu2, URA3/ura3, CRZ1/CRZ1-mTFP1-HIS3, HOG1-mNeptune2.5-kanMX /HOG1-mNeptune2.5-kanMX, PKC1-mRuby3-hphMX / PKC1-mRuby3-hphMX, SFP1/SFP1-mKOκ-hphMX, MSN2/MSN2-mNeonGreen-hphMX, CDC10/CDC10-mCyOFP1-natMX* |
| *OAM654* | *MATa/MATα, HIS3/his3, TRP1/trp1::ATG8p-mNeonGreen-ATG8-ATG8ter, LEU2/leu2, URA3/ura3, CDC10/CDC10-mCyOFP1-natMX* |
| *OAM671* | *MATa/MATα, HIS3/his3, TRP1/trp1, LEU2/leu2, URA3/ura3::CUP1p-QN-mRuby3, CDC10/CDC10-mCyOFP1-natMX* |
| *OAM673* | *MATa/MATα, HIS3/his3, TRP1/trp1, LEU2/leu2, URA3/ura3, TOM70/TOM70-mTFP1-HIS3, CDC10/CDC10-mCyOFP1-natMX* |
| *OAM670* | *MATa/MATα, HIS3/his3, TRP1/trp1, LEU2/leu2, URA3/ura3, RAD52/RAD52-mNeonGreen-hphMX, CDC10/CDC10-mCyOFP1-natMX.* |
| *OAM454* | *MATa/MATα, HIS3/his3, TRP1/trp1, LEU2/leu2, ura3 /URA3, RTG1-mNeptune2.5-kanMX /RTG1-mNeptune2.5-kanMX, SFP1-mScarlet-I-hphMX /SFP1-mScarlet-I-hphMX, GLN3/GLN3-mKOκ-hphMX, XBP1/XBP1-mNeonGreen-hphMX, CDC10/CDC10-mCyOFP1-natMX* |
| *OAM458* | *HIS3/his3, TRP1/trp1, LEU2/leu2, ura3 /ura3::CUP1p-XBP1-mTFP1-URA3, RTG1-mNeptune2.5-kanMX /RTG1-mNeptune2.5-kanMX, SFP1-mScarlet-I-hphMX /SFP1-mScarlet-I-hphMX, GLN3/GLN3-mKOκ-hphMX, XBP1/XBP1-mNeonGreen-hphMX, CDC10/CDC10-mCyOFP1-natMX* |
| *OAM466* | *MATa/MATα, HIS3/his3, TRP1/trp1, LEU2/leu2, ura3 /ura3::CUP1p-xbp1-EE-mTFP1-URA3, RTG1-mNeptune2.5-kanMX /RTG1-mNeptune2.5-kanMX, SFP1-mScarlet-I-hphMX /SFP1-mScarlet-I-hphMX, GLN3/GLN3-mKOκ-hphMX, XBP1/XBP1-mNeonGreen-hphMX, CDC10/CDC10-mCyOFP1-natMX* |
| *OAM497* | *MATa/MATα, HIS3/his3, TRP1/trp1, LEU2/leu2, ura3 /ura3::CUP1p-STB3-mTFP1-URA3, RTG1-mNeptune2.5-kanMX /RTG1-mNeptune2.5-kanMX, SFP1-mScarlet-I-hphMX /SFP1-mScarlet-I-hphMX, GLN3/GLN3-mKOκ-hphMX, XBP1/XBP1-mNeonGreen-hphMX, CDC10/CDC10-mCyOFP1-natMX* |
| *OAM455* | *MATa/MATα, HIS3/his3, TRP1/trp1, LEU2/leu2, ura3 /ura3::CUP1p-WHI5-mTFP1-URA3, RTG1-mNeptune2.5-kanMX /RTG1-mNeptune2.5-kanMX, SFP1-mScarlet-I-hphMX /SFP1-mScarlet-I-hphMX, GLN3/GLN3-mKOκ-hphMX, XBP1/XBP1-mNeonGreen-hphMX, CDC10/CDC10-mCyOFP1-natMX* |
| *OAM500* | *MATa/MATα, HIS3/his3, TRP1/trp1, LEU2/leu2, URA3/ura3, STB3/STB3-mTFP1-HIS3, RTG1-mNeptune2.5-kanMX /RTG1-mNeptune2.5-kanMX, SFP1-mScarlet-I-hphMX /SFP1-mScarlet-I-hphMX, GLN3/GLN3-mKOκ-hphMX, MSN2/MSN2-mNeonGreen-hphMX, CDC10/CDC10-mCyOFP1-natMX, xbp1Δ::kanMX4/ xbp1Δ::kanMX4* |
| *OAM502* | *MATa/MATα, HIS3/his3, TRP1/trp1, LEU2/leu2, URA3/ura3, STB3/STB3-mTFP1HIS3, RTG1-mNeptune2.5-kanMX /RTG1-mNeptune2.5-kanMX, SFP1-mScarlet-I-hphMX /SFP1-mScarlet-I-hphMX, GLN3/GLN3-mKOκ-hphMX, MSN2/MSN2-mNeonGreen-hphMX, CDC10/CDC10-mCyOFP1-natMX* |
| *OAM849 (TK52)* | *MATa/MATα, HIS3/his3, TRP1/trp1, LEU2/leu2, URA3/ura3, XBP1/XBP1-mScarlet-I-hphMX* |
| *OAM850 (PK1273)* | *MATa/MATα, HIS3/his3, trp1/trp1, LEU2/leu2, URA3/ura3, WHI5/whi5::WHI5pr-WHI5-mKOk-TRP1* |

**Supplementary Table 4. Optics setup for a six-color imaging system to study the proliferation-quiescence transition.** Excepting HE61 (Zeiss) and ET645/75 (Chroma), all other filters are from Semrock. A LED Colibri2 light source was used, setting 470, 505, 540-580 LEDs to 15% intensity and the 615 LED to 25% intensity. The combination of filters and dichroic mirrors was designed according to theoretical excitation-emission spectra using Semrock's SearchLight™ spectra viewer tool (https://searchlight.semrock.com/) and validated experimentally using strains tagged with single fluorophores to confirm the expected properties of each channel under microfluidic conditions *in vivo* (See figure S2).

| LED light source | Fluorophore | Excitation filters | Dichroic mirrors | Emission filters | Remarks |
|---|---|---|---|---|---|
| 470 | mTFP1 | FF01-445/45-25, FF01-482/35-25 | FF482-Di01-25x36 | FF01-488/6-25 | Notice how the two excitation filters create a narrow bandpass filter to reduce exposure to shorter phototoxic wavelengths. |
| 505 | mNeonGreen/ yemNeonGreeen | FF01-504/12-25 | FF518-Di01-25x36 | FF01-530/11-25 | The high quantum yield of mNG restricts its use to low abundance proteins or weak signals. Tagging abundant proteins with mNG in this system results in intolerable spillover in adjacent channels. |
| 540-580 | mKO*κ* | FF01-534/20-25 | FF552-Di02-25x36 | FF01-563/9-25 | This channel can be contaminated by the mCyOFP1 signal depending on the intensity of the protein being detected in the mCyOFP1 channel. |
| 540-580 | mRuby3/ mScarlet-I | FF01-563/9-25 | FF573-Di01-25x36 | FF01-598/25-25 | Using Scarlet-I or mRuby3 depends on the abundance of the protein to be fluorescently tagged. Scarlet-I was used for less abundant proteins or weak signals. |
| 615 | mNeptune2.5 | FF01-615/45-25, FF01-629/56-25 | FF652-Di01-25x36 | FF01-708/75-25 | Two excitation filters create a narrower bandpass filter that excludes the potential bleed-through from red fluorophores (mScarlet-I, mRuby3) |
| 505 | CyOFP1 | HE61 | HE61 | ET645/75 | As a long Stokes shift fluorophore, mCyOFP1 is compatible for imaging together with green and red fluorophores. However, strong signals in CyOFP1 can produce bleed-through in the mKO*κ* channel. |
